# Mixtures are more salient stimuli in olfaction

**DOI:** 10.1101/163238

**Authors:** Ho Ka Chan, Thomas Nowotny

## Abstract

In their natural environment, animals often encounter complex mixtures of odours. It is an open question whether and how responses to complex mixtures of multi-component odours differ from those to simpler mixtures or single components. To approach this question, we built a full-size model of the early olfactory system of honeybees, which predicts responses to both single odorants and mixtures. The model is designed so that olfactory response patterns conform to the statistics derived from experimental data for a variety of their properties. It also takes into account several biophysical processes at a minimal level, including processes of chemical binding and activation in receptors, and spike generation and transmission in the antennal lobe network. We verify that key findings from other experimental data, not used in building the model, are reproduced in it. In particular, we replicate the strong correlation among receptor neurons and the weaker correlation among projection neurons observed in experimental data and show that this decorrelation is predominantly due to inhibition by interneurons. By simulation and mathematical analysis of our model, we demonstrate that the chemical processes of receptor binding and activation already lead to significant differences between the responses to mixtures and those to single component stimuli. On average, the response latency of olfactory receptor neurons at low stimulus concentrations is reduced and the response patterns become less variable across concentrations as the number of odour components in the stimulus increases. These effects are preserved in the projection neurons. Our results suggest that the early olfactory system of insects may be particularly efficient in processing mixtures, which corresponds well to the observation that chemical signalling in nature predominantly involves mixtures.

## 1 Introduction

Unlike in other sensory modalities, the perceptual structure of the space of olfactory stimuli is poorly understood. Animal experiments addressing odour coding have predominantly been conducted with individual simple organic compounds (Galizia et al, 1999; Sachse and Galizia, 2003; Stopfer et al, 2003; Bhandawat et al, 2007; Carcaud et al, 2012) or few component mixtures (Rosapars et al, 2008; Deisig et al, 2010; Meyer et al, 2012; Cruz and Lowe, 2013). However, odour sources in animals’ natural environments typically emit complex mixtures of many different compounds (Knudsen, 1999; El-Sayed et al, 2008; Hetherington-Rauth et al, 2016) and signals used in chemical communication between animals are predominantly mixtures (Blum, 1992; Bortolotti and Costa, 2014). Moreover, while lab experiments on odour recognition and learning use controlled odour concentrations, odours in natural environments propagate in complex turbulent odour plumes in which filaments of high concentration are interweaved with large swathes of volume at significantly lower concentration and even clean air (Murlis et al, 1992;Weissburg et al, 2012). To advance our understanding in this area, we developed a mathematical model of the early olfactory system of bees and investigated whether and how sensing individual compounds may differ from sensing mixtures.

A number of computational models of the bee olfactory system have been proposed (Hopfield, 1991,1995; Rospars et al 2008; Luo et al, 2010; Nowotny et al, 2013) but it has remained very difficult to combine bio-physically realistic processes and existing experimental observations into predictive mathematical models. One particular challenge is that experimental data is necessarily limited in its scope because of limitations of experimental techniques. For example, in honeybees, downstream responses of only around 30 of a total of 160 identifiable olfactory receptor neuron (ORN) types can be routinely measured (Galizia et al, 1999; Sachse and Galizia, 1999). When constructing a data-driven model based on these measurements, Nowotny et al (2013) limited their model of the bee olfactory system to the 30 ORN types for which data is available. However, synaptic efficacies, connection patterns and sources of noise cannot be scaled to generate a fully faithful mapping from 160 to 30 ORN types. In addition, when creating response patterns by directly fitting experimental data on a bespoke subset of ORN types (Nowotny et al., 2013, Rospars et al, 2008), it cannot be expected that the resulting predictions would reflect the properties of other parts of the system outside the scope of the original experimental measurements. Because of this and large animal-to-animal variability in olfactory responses, it would be unlikely that the model was consistent with other experimental data not used for building it. Other models use more phenomenological approaches, for example by employing random patterns as inputs (Hopfield, 1991, 1995; Luo et al, 2010, Huerta et al., 2004, Nowotny et al., 2005). This can provide important insights into the putative role of different elements of the neural circuits but is unlikely to offer direct, experimentally testable predictions.

Here we present a third, alternative approach, where the model is designed to consider inputs representative of the full sensory input space of the animal as well as relevant biophysical processes. To be specific, we have built a model of the honeybee olfactory system, which comprises a full set of 160 ORN types and corresponding olfactory glomeruli in the antennal lobe (AL). In the ORNs, the model accounts for the processes of odour binding and receptor activation, as well as for the non-linearity of spike generation. ORN activity then excites local neurons (LNs) and projection neurons (PNs) in the AL, which interact with each other to generate the output activity of PNs. In order to relate to known experimental results, the response characteristics of ORNs in the model were tuned to match the statistics of olfactory responses in bees from a number of existing data sets (see section 2 for details), but not to any particular individual measurements. While this does not allow to make predictions for responses to specific chemical compounds, this approach is the most likely to yield meaningful predictions in a‘typical’, or,‘on average’ sense. Crucially, using in addition a biophysical model for binding and activation allows us to predict both, responses to individual compounds and to arbitrary mixtures of such compounds.

In addition to the statistics it was built to reflect, our model also reproduces key features of ORN and PN responses observed in separate experimental work that was not considered when building the model. It replicates the pulse tracking ability of ORNs (Syzszka et al, 2014) (Section 3.1.1) and the wide range of dose-response relationship in PNs (Ditzen, 2005) (Section 3.1.2). Furthermore, we were able to identify biophysical processes which may contribute to the decorrelation among PN responses across different glomeruli (Section 3.1.3) (Ditzen, 2005) and to observed statistical differences between ORN and PN responses (Section 3.1.4) (Deisig et al, 2010; Luo et al, 2010).

We analysed the receptor dynamics for mixtures and single component odour stimuli. The model reproduces both synergistic and hypoadditive responses as observed experimentally (Duchamp-Viret et al, 2003; Rospars et al, 2008; Cruz and Lowe, 2013) (Section 3.2.1). On the systems level, the model predicts that both the asymptotic (Section 3.2.2) and transient responses (Section 3.2.4) are larger for mixture stimuli than for simple odours, when the stimulus concentration is small. The latter also implies a smaller ORN response latency for mixtures than for simple odours in this regime. In addition, the model predicts that the response gain at small concentrations is more correlated to the asymptotic response at large concentrations for mixture stimuli than for simple odours, which implies more concentration-invariant ORN response patterns for mixtures (Section 3.2.3). These predictions were obtained both analytically and numerically and extend beyond ORNs to the PNs in the AL. Our results suggest that the olfactory system may be more efficient and reliable in coding mixtures than simple odorants.

## 2. Methods

### 2.1 Model network

Our model describes the early stages of the olfactory system of the honey bee, consisting of ORNs on the antenna, and PNs and LNs in the AL (see Figure 1). ORNs of the same type respond according to the same response profile and connect to the same glomerulus in the AL. Here they synapse on LNs and PNs and the PNs relay the processed olfactory information to higher brain centres such as the mushroom bodies and parts of the lateral protocerebrum. LNs are local to the antennal lobe and modify the PN response pattern through inhibition (Figure 1).

**Figure 1:**
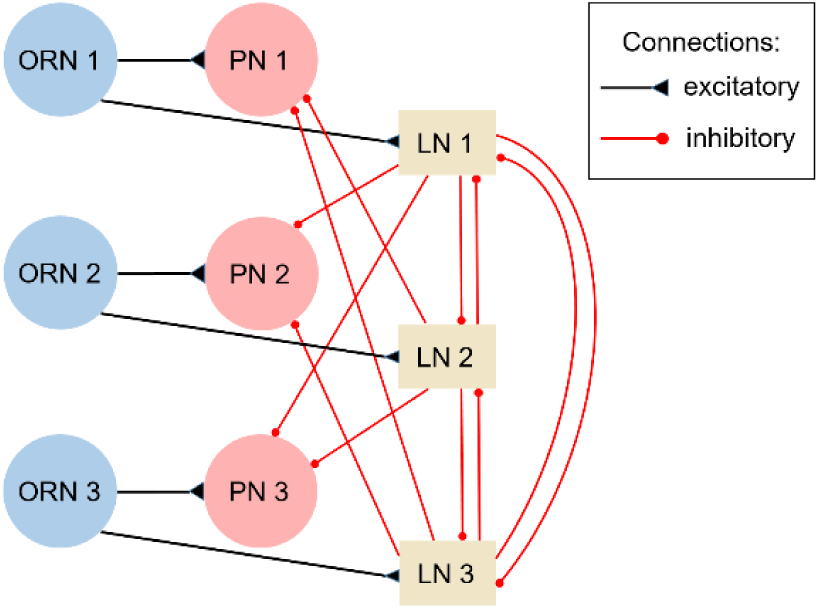
Illustration of part of the model AL network. ORNs excite LNs and PNs of‘their’ glomerulus, and LNs then project with inhibitory synapses to the PNs and LNs of other glomeruli.

In our model, responses from the same type of ORNs, LNs or PNs are approximated by their ensemble average. We therefore use a single unit to represent all units of the same type. In this work, we refer to each type of a certain entity by its representatives (e.g. 20 types of ORNs will be referred to as 20 ORNs). Please refer to Appendix A for the parameters used in the model.

### 2.2 Generating ORN responses

The olfactory response begins with the transduction of volatile chemicals in the air into membrane conductance changes of ORNs. We construct the model to reproduce the statistical properties of fully temporally and concentration resolved ORN responses in a number of steps. First, we consider the asymptotic response of model ORNs for time-invariant odour inputs at high concentrations.

#### 2.2.1 Asymptotic response of model ORNs at high concentrations

Asymptotic responses of 28 ORNs for time-invariant odour inputs at high concentration to 16 different odours have been measured using calcium imaging of glomeruli with bath-applied Ca^2^+ dyes (Galizia et al, 1999). We adopted these responses directly to form the responses of the first 28 ORNs in our model. We then generated the responses of the remaining 132 ORNs to the same 16 odours using a method inspired by (Haenicke, 2015). The response patterns are generated from a combination of previously generated responses, including those experimentally observed in Galizia et al (1999), and noise. The ratio of the combination is determined by a target similarity matrix of odour response patterns and a global variable which determines the overall amount of correlations across the response patterns (please refer to Appendix B). The parameters are chosen such that the statistical distribution of the pairwise correlations of ORNs across odours in the generated responses matches that of the 28 ORNs adopted from data. The generated responses are then rescaled such that the mean and the variance of the responses for all receptor-odour combinations as well as the mean and the variance of the variance of the responses across odours for each ORN match. Please refer to the Appendix A for the detailed method how this is achieved.

ORN responses to chemically similar odours are correlated (Carcaud et al, 2012). In our model, such correlations are quantified using the normalized Euclidean distances *d*_*ij*_ between the response vectors of 2 different odours *i* and *j*, denoted by *x*_*i*_ and *x*_*j*_, as in Carcaud et al (2012).

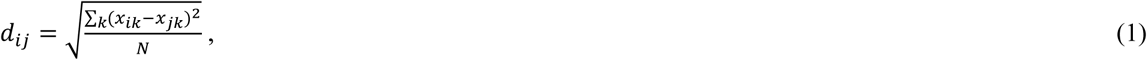

where *N* is the total number of odours in our input space and the subscript *k* labels the different ORNs.

The responses of all previously generated ORNs are then iteratively tuned so that the Euclidean distance matrix d for the generated response patterns matches the distance matrix observed in the experimental data. The tuning processes are designed to limit changes to the statistical quantities calibrated previously.

#### 2.2.2 Asymptotic ORN response to odours at other concentrations

We model the dose-response relationship by Hill curves.

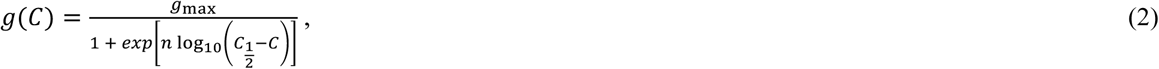

where *g* is the ORN response, and *C* = log(*c*) is the logarithm of the concentration *c*. The Hill coefficient *n* and the inflection point *C*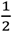 are sampled from log-normal and normal distributions respectively as experimentally observed (Gremiaux et al, 2012). The asymptotic maxima are chosen such that the responses to odours at concentration c =1, corresponding to undiluted stimuli, match the asymptotic responses described in section 2.2.1 above.

#### 2.2.3 Temporally resolved ORN response

The time series response to a single odour stimulus is generated using a set of ordinary differential equations describing the binding and activation of receptors as in Nowotny et al (2013):

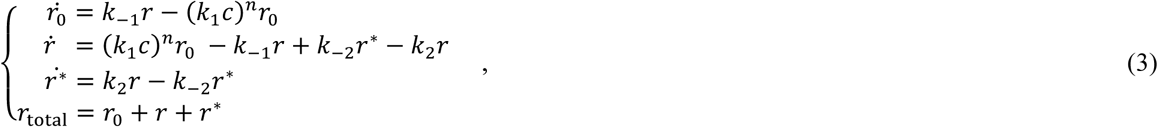

where *k*_1_(*k*_‒1_) and *k*_2_(*k*_‒2_) are the (un)binding constants and (de)activation constants respectively, *c* is the concentration of the odour, *n* describes the effects of the transduction cascade, and *r*_0_, *r* and *r*_i_* are the‘effective concentration’ of free, bound and activated receptors such that *r*\* is proportional to an excitatory conductance of the ORN (see 2.4 for more details). The sum of the number of receptors in different states is equal to the total number of available receptors *r*_total_, as described by the last equation.

The response-dose relationship at large t can be described by (2) when a time-invariant stimulus is applied (Rospars et al 2008; see also Appendix C). *n, k*_1_(*k*_‒1_) and *k*_2_(*k*_‒2_) are partially constrained by the parameters in the Hill curves for the asymptotic responses as described in section 2.2.2. In dealing with the remaining degrees of freedom, we take into account the typical timescale of dynamics in AL responses measured experimentally (Syzszka et al, 2012, 2014). Note that while k1, k_‒1_, k2 and k_‒2_ depend on both the receptor types and odorants, *n* only depends on receptor types, independent of odorants.

When a mixture of individual odour compounds is present, the time series response can be described by a slightly modified version of (3).

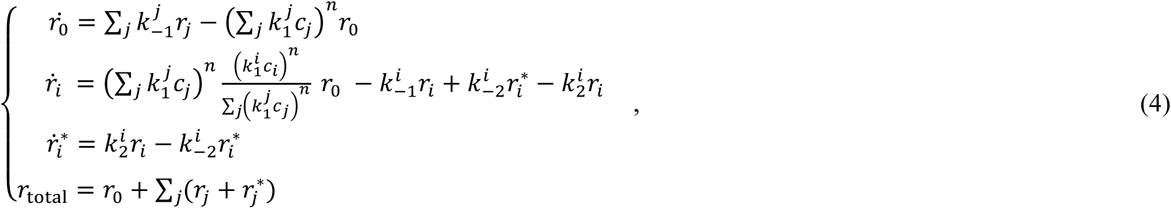

where the subscript i indicates that the corresponding quantities describe the ith component in the mixture. Unlike similar models in previous work (Rospars et al, 2008; Nowotny et al, 2013), equation (4) does not have any inconsistencies when considering‘mixtures’ of identical components with randomly partitioned concentrations. Apart from Section 3.2.5 and Appendix C, we only consider a scenario where the concentrations of all the constituent compounds in a mixture are identical, i.e. c_1_ = c_2_ =… = c. The subsequent analysis is based on this assumption.

The asymptotic solution for (3) and (4) can be obtained analytically (See appendix C). We have also simulated the equations directly, integrating (4) numerically for 250ms for each odour-receptors combination. At the end of the simulation, the receptors converge extremely well to the asymptotic responses determined analytically (results not shown).

### 2.3 Generating PN responses

In our model, ORNs provide excitatory input to PNs and LNs, all ORNs of a given type to the same glomerulus. PNs also receive inhibitory input from the LNs of all other glomeruli. This is illustrated in Figure 1.

To be consistent with the findings in Linster et al (2005), for any pair of glomeruli *i* and *j*, the connections from the the LN in glomerulus *j* to the PN and LN in glomerulus i have weight *W*_*ij*_, which is a function of the correlations *ρ*_*ij*_ between the corresponding ORN response patterns,

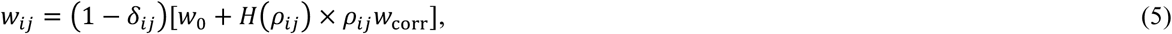

where *δ*_*ij*_ is the Kronecker delta, *H* is the Heaviside step function, *W*_0_ and *W*_corr_ are normal distributed random variables, and *ρ*_*ij*_ is the Pearson correlation between the conductance of ORN i and j when stimulated by odours at high concentration, as obtained in 2.2.1.

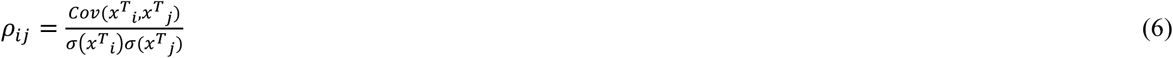

In section 3.1.3, we will also show results for the two additional cases where (i) all LN-PN connections are removed and (ii) where the strength of LN-PN connections are uniform instead. Please note that in the model, PNs do not receive feedback inhibition from LNs in the same glomerulus since such connections, should they exist, can be effectively taken into account to by tuning the ORN-PN excitation.

### 2.4 Obtaining the instantaneous firing rate of neurons

The responses we generated in the previous part are assumed to describe conductances. To obtain the firing rate of the ORNs from its input conductances, we approximated the dynamics of a neuron by the conductance-based leaky integrate-and-fire model with adaptation (Richardson and Gerstner, 2005; Chan et al., 2016).

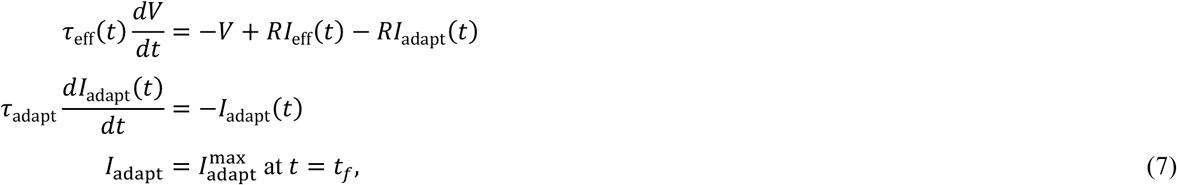

where *V* is the membrane potential, *R* is the membrane resistance, and I_adapt_ is the adaptation current, which is set to 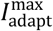 just after firing events at *t*_*f*_ and decays exponentially with decay time constant *τ*_adapt_. *I*_eff_ and *τ*_eff_ are the effective input current and effective membrane time constant having taken into account the conductance effects. They are described by

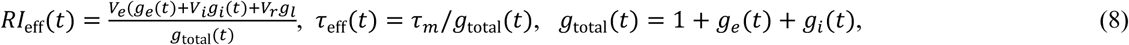

where *V*_r_ is the membrane rest potential, *V*_*e*_ and *V*_*i*_ are the reversal potentials of excitatory and inhibitory synapses, *g*_*l*_ is the membrane leak conductance, *g*total(*t*) is the total conductance of the neuron, ge and gi are the excitatory and inhibtory conductance. For ORNs, 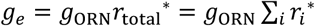, where *g*_ORN_ is a constant, and *g*_*i*_ = 0. For PNs and LNs, *g*_*e*_ and *g*_*i*_, as a first order approximation, are proportional to the firing rate of ORNs and LNs (Kuhn et al, 2004). For PNs, we also add constant background noise into *g*_*e*_(*t*) and *g*_*i*_(*t*) so that they fire spontaneously at 5 - 20Hz. We then set R = 1 by absorbing it into 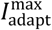 and other variables. When *V* reaches the threshold Vth, the neuron fires a spike and *V* is immediately reset to Vreset.

We then adopt the adiabatic approximation by considering the input to be quasi-time-invariant on the time scale of neuronal firing, such that *τ*_eff_(*t*) and *I*_eff_(t) are taken to be constant. With the additional assumption of noise-free input and setting *t*_*f*_ = 0, the membrane potential before the next firing event can be obtained analytically as follows:

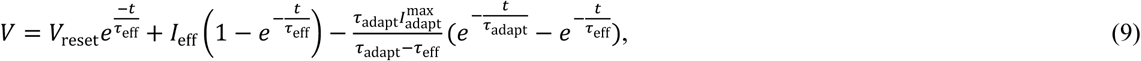

where *V*_reset_ is the reset potential after the neuron has fired. The instantaneous firing rate of the neuron can then be obtained using:

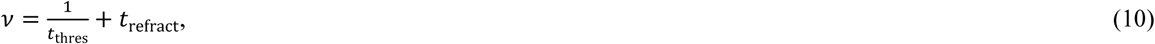

where *t*_thres_ is the time when *V* = *V*_th_, which is to be obtained numerically, and trefract is the absolute refractory period. We may then use (9) and (10) to calculate v at different time points for fluctuating input. Please note that *v*, by our definition, does not directly relate to temporal information of spike patterns, e.g. the inter-spike interval between any pair of spikes (see discussion).

The calculations for LN and PN responses are iterated several times to allow the system to settle into a steady state. Here we ignore any oscillation or other complex dynamics that may take place in biological ALs.

In this work, we take 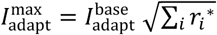 for ORNs, whe and 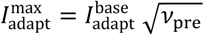 for PNs and LNs, where 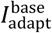 is a constant and vpre is the firing rate of the corresponding units in the previous iteration. However, qualitatively similar results can be obtained by assuming 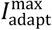 to be constant (results not shown).

### 2.5 Obtaining the response latency of ORNs

As we have discussed in the last session, the response latency of neurons, defined by the time taken for the neuron to fire the first spike after stimulus onset, cannot be obtained from the instantaneous firing rate. Instead, we directly integrate (7) numerically, assuming *V* = *V*_mean_ at *t* = 0 and obtain the response latency by finding tthres. While we consider time-invariant constant noise (with the main purpose of bringing the steady state of *V* closer to the firing threshold and increasing its responsiveness to inputs), we still have not incorporate any element of stochasticity here and therefore cannot account for noise-driven responses. Nevertheless, the adiabatic approximation is not used here. Therefore the results remain valid for strongly fluctuating input provided the neural response is mean-driven.

## 3.Results

### 3.1 Comparison between model results and other published experimental data

#### 3.1.1 Mean ORN response to pulse input

It has been shown that ORNs of honey bees can track pulses of odours up to high frequencies on the order of 100Hz (Szyszka et al, 2014). When they are repeatedly stimulated by short odour pulses, their average responses, as measured by the electro-antennogram, exhibit oscillations with the same frequency as the odour pulses, defined as the reciprocal of the inter-pulse interval. The amplitude of the oscillations decreases with the pulse frequency (See Figure 2a).

**Figure 2:**
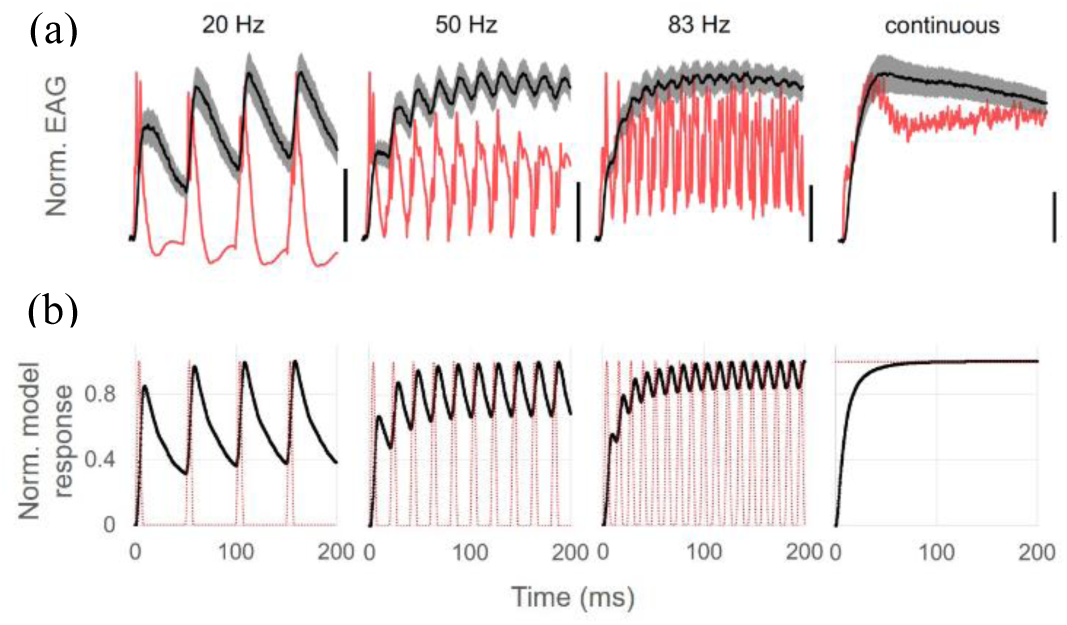
ORN responses to pulsed and constant stimuli as measured by electro-antennogram recordings Syzszka et al (2014) are qualitative similar to the average normalized ORN responses to the corresponding stimuli (1-hexanol at *c* = 10-^4^) predicted by our model (b). The red lines show the concentration of the odour input pulses.

We stimulated our model ORNs with short odour pulses of ∼5ms duration at different frequencies. Figure 2b shows that the average responses and their frequency-dependence are qualitatively similar to experimental measurements as shown in Figure 2a. The model neurons show slightly better pulse tracking capability than their experimental counterparts, likely due to the lack of a temporal filter for neural firing under the adiabatic approximation (see 2.4) (Ostojic and Brunel, 2011).

#### 3.1.2 Asymptotic response-dose relationship of PNs

While the asymptotic ORN responses are almost always monotonically increasing with the concentration of the stimulus (Sachse and Galizia, 2003; Carcaud et al, 2012), it has been widely reported that this is not always the case for PNs (Sachse and Galizia, 2003; Yamagata et al, 2009). Experimental data in Ditzen (2005) illustrates that a wide range of individual response-dose relationships exists for PNs, which we can summarize as follows: The response monotonically increases/decreases with stimulus concentration (increase/decrease); the unit is inactivated for all concentrations (inactivated); the response has a non-monotonic relationship with concentration (other). The fraction of occurrence of each type of relationship is shown in Figure 3.

**Figure 3:**
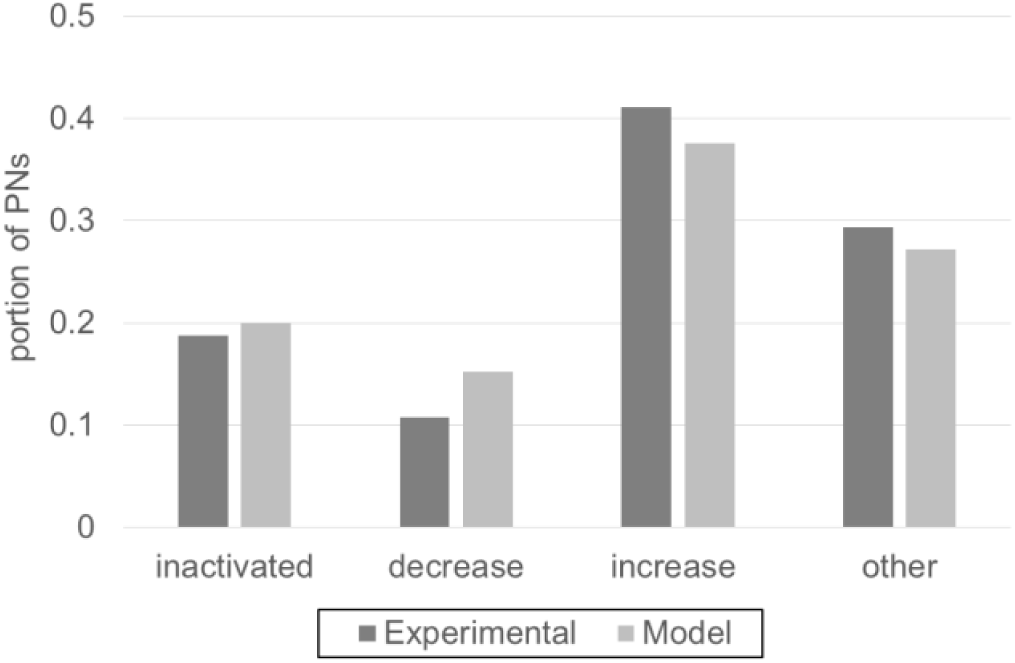
Statistical distribution of different kinds of response-dose relationships observed in model PN responses (light grey bars) and experimental PN data from Ditzen (2005) (dark grey bars). We divide response-dose relationships into 4 different types: “inactivated” where PNs show no or very weak responses (<16Hz) to stimuli at any dose, “decrease” (“increase”) where responses decrease (increase) with dose and “other” where responses are independent of or display non-monotonic relationships with dose. All types of response-dose relationships can be generated by our model. With appropriate choice of parameter, the portions of each type generated by our model can match those observed in experimental data.

Our model can reproduce all observed types of relationships, as shown in Figure 2. Note that the parameters of the model have been slightly tuned to match the proportion, with which each type of relationship occurred to the experimental data, but all the relationships can be reproduced within a large area of parameter space. The non-monotonic relationships are produced as the result of LN inhibition in the AL network. Without inhibition from LNs, only the‘increase’ and‘inactivated’ type relationship can be recovered (results not shown).

The presence of cases of decreased in responses with increasing stimulus concentration leads to a weaker scaling of the mean PN activity with concentration (See Figure 6), in particular in the high concentration regime, where inhibition by LNs is strong.

**Figure 6:**
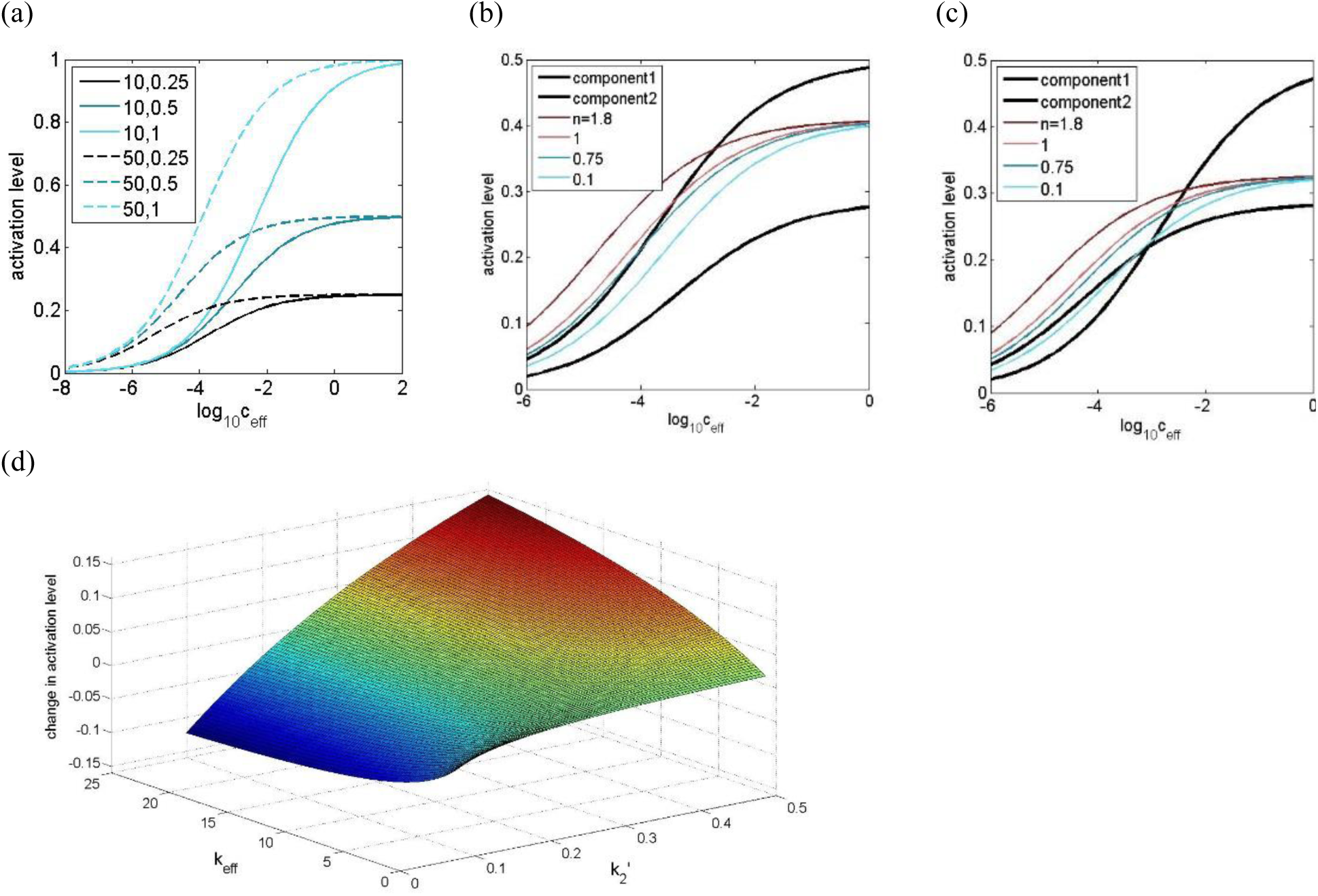
(a) The activation level of receptors for which the dynamics are described by (3). In the limit of small ceff, receptors with the same Keff respond identically, regardless of the value of 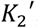. In the other limit of large ceff, responses always approach an asymptotic value depending on 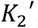, regardless of the value of Keff. Legend format: 1st number: Keff, 2nd number:, 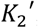. (b,c) Examples of the response to binary mixtures as *n* varies. The responses to their constituent components are shown as thick black lines. In the limit of small ceff, the mixture response can be synergistic, linearly additive and hypoadditive depending on the values of n. In the other limit, the responses to mixtures are independent of the value of *n* and are always in between those of their constituent components. For simplicity, we chose k1 for both components to be the same.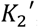, Keff for each component in (b): 0.29,8.33;0.5,20 (c): 0.5,8.33;0.29,20. (d) The change of activation level of a receptor when another component is added to the original single component stimulus. At high 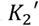 (for the added odour), the response to the added odour alone, and therefore the weighted mean of the response, is higher than that of the original odour, resulting in a rise in response after the addition of the odour, and vice versa. The larger the Keff (for the added odour), the larger the weight for the response of the added odour, resulting in a larger deviation of the response from that of the original odour. 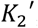, Keff for the original odour: 0.25,10.

#### 3.1.3 Correlation between asymptotic PN responses

It has been shown that while the asymptotic ORN responses across odours are highly correlated (Galizia et al, 1999, see also Figure 4a), correlations of PN response patterns are in general much weaker (Ditzen, 2005, see also Figure 4a). Our model is designed to match the correlations of ORN responses observed in experimental data, as described in 2.2.1. It is of interest to study how well the correlations in generated PN responses match their experimental counterpart, which was not directly fitted to in the construction of the model. Figure 4b shows the probability distribution of pairwise correlations between experimental and model ORN and PN responses across odours. The model ORN responses are, as expected, highly correlated, but the degree of correlations is slightly less than that observed in the experimental results. This is likely a result of tuning for chemical similarity (See section 2.2.1) and non-linearity in spike generation. On the other hand, the distributions for PNs also exhibit a good qualitative match with its experimental counterpart, as both have a peak around correlation = 0. Please note that in Figure 4b, the model responses are in the form of firing rates while the experimental responses are obtained from calcium imaging data, which is strongly related to the former but may not be directly proportional to it (Grienberger and Konnerth, 2012).

**Figure 4:**
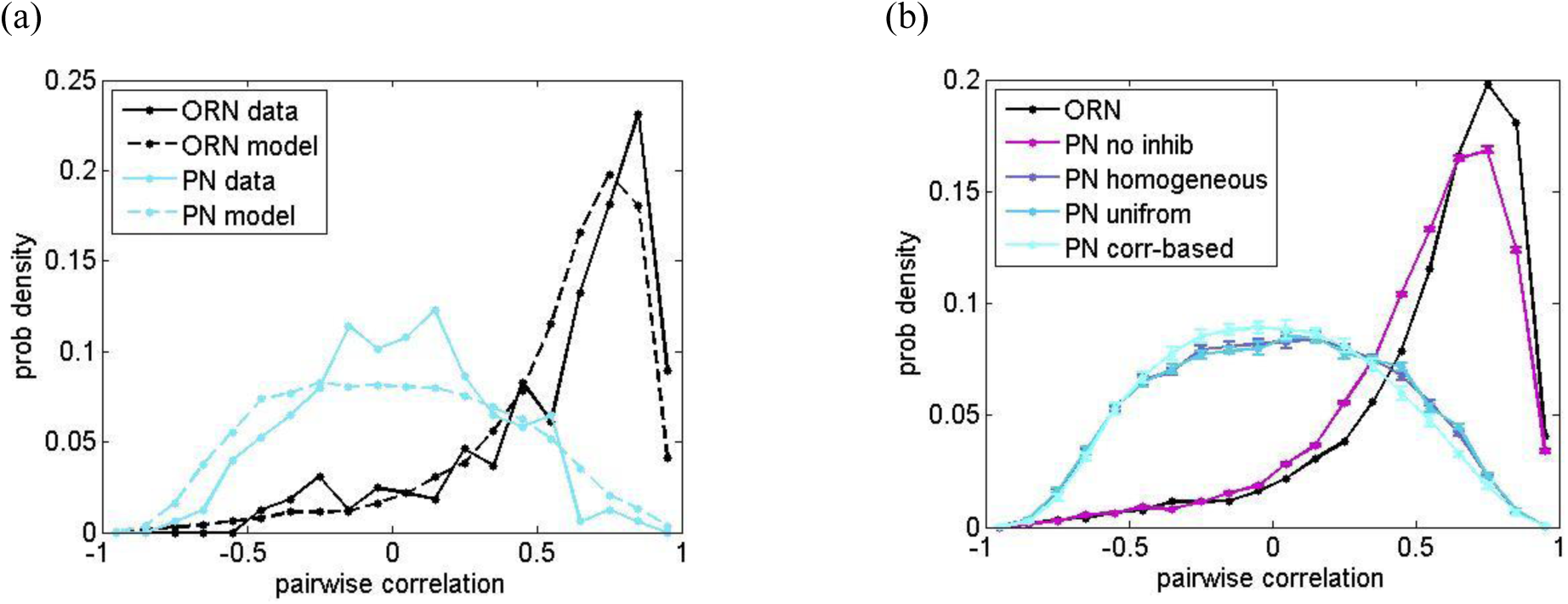
(a) Statistical distribution for pairwise correlation across response patterns for ORNs (concentration c=1) and PNs (c=0.1) observed in calcium imaging experiments (Galizia et al, 1999; Ditzen, 2005) and our model. Our methods are designed to fit the distribution of correlation across ORNs observed in experimental data when building the model. No such fitting is done for the PNs. Please also note that in the plot, the‘model responses’ correspond to the firing rate of units which may not be directly related to the response obtained by calcium imaging. (b) Comparison between pairwise correlation across firing rate patterns for ORN and PN with no inhibition (purple), inhibition with homogeneous (blue), uniform and normal distributed (green) and correlation-based (light green) connectivity. The parameters for the latter three scenarios are chosen such that the mean output firing rate of PNs are the same for all 3 cases. While the non-linearity in neural spiking and the specific correlation-based PN-LN connectivity patterns contributed a little to the decorrelation, the most significant contributing factor is the presence of inhibition. The results for PNs are obtained over 10 trials from the same set of ORN data and the error bars correspond to standard deviation across trials. c=0.1 for all the scenarios in (b).

The decorrelation of the PN response may occur in several distinct processes. Non-linearity in neuronal spiking models (de la Rocha et al, 2007; Rosenbaum and Josic, 2011) and neural inhibition can both lead to decorrelation (Middleton et al, 2012; Tetzlaff et al, 2012). To elucidate the source of the decorrelation in our model, we compared the PN responses generated from AL networks with different inhibition paradigms. Figure 4b shows that the non-linearity in the LIF model, the inhibitory neural network involving LNs and the specific correlation-based PN-LN connectivity all contributed to the decorrelation in PN activities. However, the dominant effect is the presence of inhibition. This supports the hypothesis by Olsen and Wilson (2008). The weak decorrelating effects due to non-linearity in neural spiking may possibly be due to the adaptation we introduced and that makes the f-I curve of the LIF neuron more linear (Ermentrout, 1998). On the other hand, the effect of correlation based PN-LN connectivity is mainly reducing the amount of strongly correlated PNs by introducing strong mutual inhibition between them. Finally, we looked at the case when all the connections between the same unit type in the AL network are homogeneous (i.e. removing variability for all connection strengths). The results exhibit no major qualitative difference to the results where the LN-PN connections are uniformly distributed, suggesting that the heterogeneity in connectivity plays only a very minor, if not negligible, role in shaping the correlation across PNs.

#### 3.1.4 Comparison between ORN and PN response

Experimental measurements show that the correlation between receptor neuron and AL activity is around 0.6-0.7 (Deisig et al, 2010). We calculate the correlation between our model ORNs and PNs odour by odour, and it shows a good match to experimental data. When LN inhibition is removed, the correlation increases but still substantially smaller than unity. This suggests that both LN inhibition and non-linearity of LIF model contribute to the differences between ORN and PN response patterns.

We next investigate the phenomenon of‘magnitude equalization’ in PN responses proposed by Luo et al (2010). In their work, the authors produce a model in which the mean of PN responses across glomeruli is less variable for different odours than the mean of ORN responses. Here, we quantified this by the coefficient of variation (CV) of the mean firing rate across glomeruli for different odours. Table 1 shows a drop in CV from the ORNs to the PNs, confirming the effect of PN‘magnitude equalization’.

**Table 1:** The mean firing rate for ORNs, PNs in the full model and PNs without inhibition. All results are averaged over 10 trials. Entries format: mean ± standard deviation across the 10 trials

|  | mean | sd | CV |
| --- | --- | --- | --- |
| ORN | 67.41 | 20.11 | 0.298 |
| PN with inhib | 40.19±1.04 | 2.42±0.15 | 0.059±0.004 |
| PN no inhib | 290.95±1.67 | 46.84±0.26 | 0.161±0.001 |

As in the last section, we compared the above results to the case where there is no inhibition. Under this circumstance, the ORN-PN correlation is significantly higher but still considerably smaller than unity (see Figure 5) while‘magnitude equalization’ is weaker but clearly significant (see Table 1), suggesting that both the non-linearity in the LIF model and LN inhibition are important in producing both phenomena.

**Figure 5:**
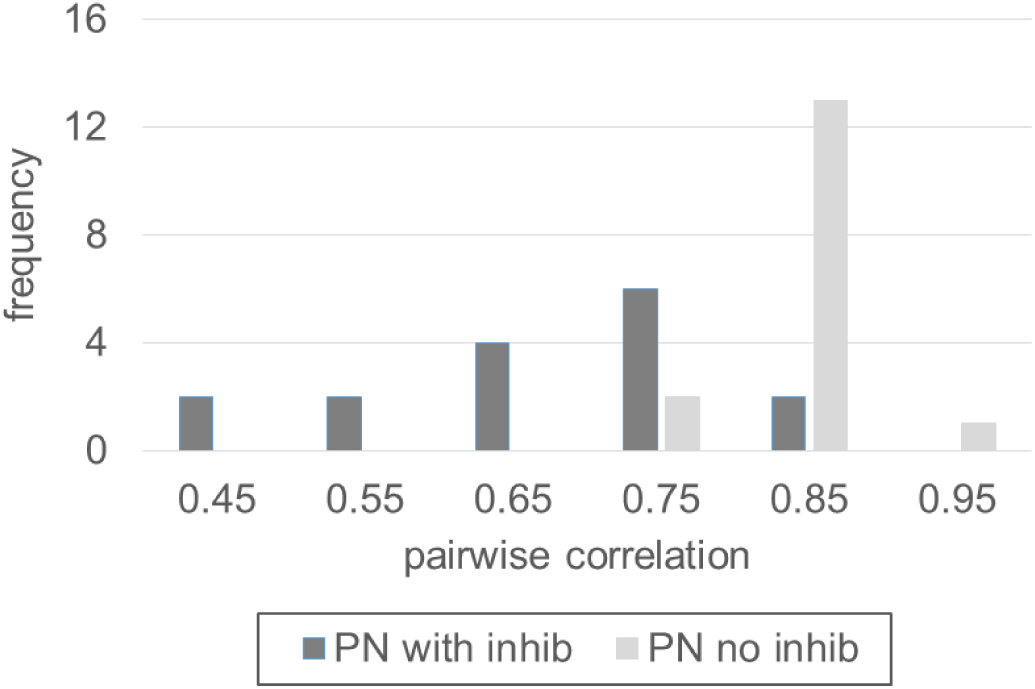
Statistical distribution of the pairwise correlation between the overall ORN and PN response for different odour stimuli. The average correlation is around 0.6-0.7, which matches well with experimental observations. Note that if we remove the inhibitory input from LNs, the correlation becomes higher but is still significantly below unity, which suggests that both, the non-linearity in the LIF model and LN inhibition contribute to differences between ORN and PN response patterns.

Please note that the above-mentioned effects may be further amplified by noise in the system. While we do add external noise to ORNs, for the purpose of obtaining realistic response latencies as described in the next section, and PNs, for the purpose of mimicking their spontaneous activity, the noise effects are small and only visible at low stimulus concentrations. The general effects of noise on neural response and correlation have been well studied (Shadlen and Newsome, 1998; Brunel et al, 2001; Ostojic et al, 2009; Chan, 2015; Chan et al, 2016) and are beyond the scope of this study.

### 3.2 Responses to mixtures

In this section, we compare the response to a mixture with that to a single component odour, as predicted by our model. We also show results from a mathematical analysis which can explain observations regarding the similarity and differences between the response patterns in these two cases. Since the only differences between mixtures and single component odours manifest themselves at the level of receptors, the analysis here is focused on receptor dynamics. The results below are obtained under the simplifying assumption that the concentrations of the odour components in mixtures are all equal, as well as equal to the concentration of the single component odours to which we are comparing. Please refer to Appendix C for the general case of heterogeneous concentrations among components in a mixture.

#### 3.2.1 Asymptotic response to mixtures and the role of the non-linear transduction process

In order to understand whether and how the receptor dynamics described by (3) and (4) may lead to qualitative differences between responses to single component odours and mixtures, we compare the results for single component odours and for mixtures in which all components have the same concentration c. We can solve the equations for r* and *r*_mix_* in (3) and (4) at equilibrium analytically, leading to (see Appendix C for the derivation):

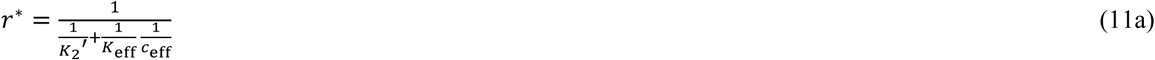

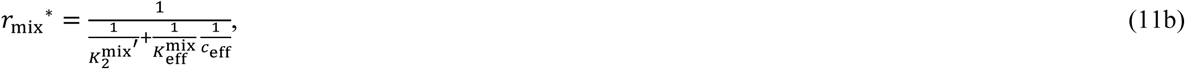

Where 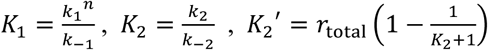, 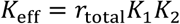, 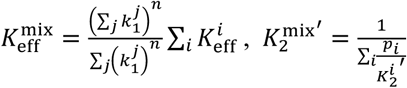, 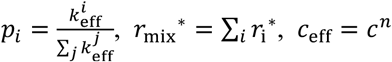 and 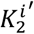 and 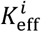 refer to the value of 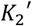 and Keff for the *i*th odour component in the mixture stimulus.

We now study the response to stimuli in the limit of low and high concentrations. It is clear from (11a) and (11b) that the responses are determined by Keff (or 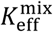 for mixtures) and 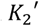 (or 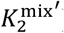). This is illustrated in Figure 6a. In the limit of small ceff, r* and *r*_mix_* are given by

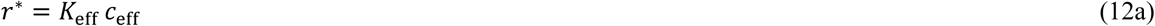

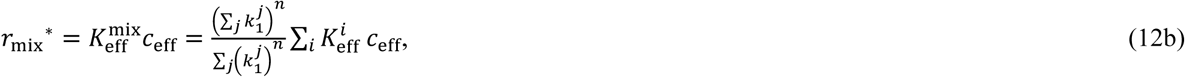

Rospars et al (2008) showed that responses to mixtures can be superlinear to the sum of the components’ responses (synergy), sub-linear but stronger than the weakest component’s response (hypoadditivity) and weaker than the weakest component’s response (suppression) (Note that our definition of these terms are different from Rospars et al, 2008). In the regime of low ceff, the interaction between odour molecules is dominated by cooperative and inhibitive transduction mechanics. In the context of our model, (12b) can reproduce both hypoadditive and synergistic responses depending on the value of n.

In appendix E, we show that for mixtures with *N* components

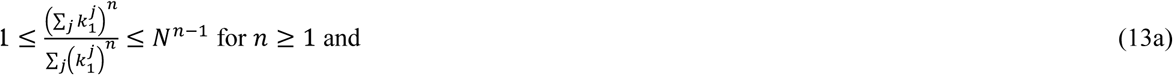

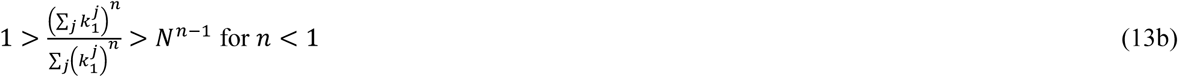

(13a) shows that synergy can be achieved when *n* > 1, hypoadditivity when 0 < *n* < 1. When *n* = 1, the responses are strictly additive. The role of *n* in mixture responses is illustrated in Figures 6b and c.

Note that even though it is possible to obtain suppressive mixture response when *n* ≤ -1 (See appendix E). We do not consider cases of non-positive n, as in such cases, the responses remain finite (when *n* = 0) or blow up (when *n* < 0) if *c* tends to 0, which is highly unrealistic.

Data analysis of experimental measurements (Gremiaux et al, 2012) shows that for biological systems, the coefficient *n* ranges takes values between 0 and 1 for most receptor types. This resonates well with other observations that responses to mixtures are predominantly hypoadditive (Duchamp-Viret et al, 2003; Rospars et al, 2008; Cruz and Lowe, 2013).

In the limit of large *c* eff, we have 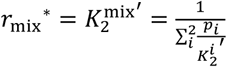 for mixtures. The response to the mixture is the weighted harmonic mean of the maximum responses to the constituent components when they are presented alone, with the

Weight 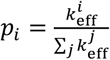 being proportional to their respective response gain at low concentrations. This implies that in this limit, the mixture response must be hypoadditive and in between the responses of its constituent components,regardless of the value of n, as shown in Figure 6b and c. This result is supported by Duchamp-Viret (2003), who showed that there are only 3% of instances where this is untrue at a high (but finite) concentration. Intuitively, in such a regime, competition for receptor sites dominates the interaction between odour molecules of different types, which gives rise to the hypoadditivity in responses. How the mixture responses are affected by 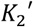 and K_eff_ of its constituent components is illustrated in Figure 6d.

#### 3.2.2 Dose-response relationship at system level

In (11a) and (11b), the parameters correspond to specific individual odour-receptor combinations. If we consider the entire space of possible odour inputs and the space of all possible chemical receptors, we would have a very large number of possible odour-reception combinations. Each combination i is characterized by parameters,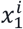,…, 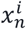, which are sampled from parameter sets X_1_,…, X_n_, each having the same number of elements as the number of possible odour-reception combinations. If we consider a sufficiently large number of such combinations, it is sensible to describe 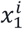,…, 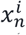, as random variables with some appropriate probability distribution each. We will take this view for all parameters in (11a) and (11b) below, which allows us to study the statistical properties of responses analytically. Note that we are not applying the above treatments to the parameter n, which reflects the properties of receptors only and is odorant-independent.

Figure 7 shows our model’s prediction of average ORN responses for single component, binary and ternary mixtures stimuli using statistically constraint parameters (See methods). At low concentration, the responses to binary and ternary mixtures are larger than those to the single components but less than twice and three times those of single components. As shown in the following, this derives from the fact that *n* < 1 for most of the receptors: Considering the average mixture response ⟨rmix*⟩,

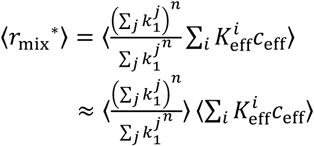

if we assume that 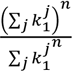 and 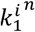 are essentially uncorrelated. While they are not truly independent, this is a good approximation since the former does not scale with 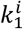 Then,

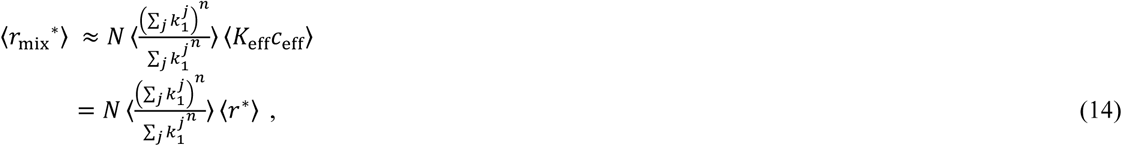

where we made use of the fact that all the 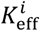 are independent and identically distributed random variables.

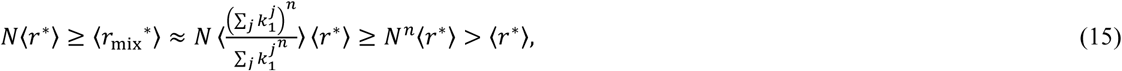

Which resonates with the findings in Figure 7. Similarly, it can be shown that N⟨r*⟩ = ⟨rmix*⟩ = Nn⟨r*⟩ if *n* = 1.

At high concentrations, our model predicts equivalent average responses to single odours and mixtures. Here we investigate the relationship between the average response to a single component odour at the limit of large C over all possible odour-receptor combinations and the average response to mixtures for the example of binary mixtures. It can be proven that for binary mixtures, ⟨r*⟩ = ⟨rmix*⟩, i.e.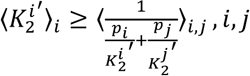 if K_eff_ is homogeneous for all odour-receptor combinations, i.e. 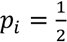, ∀i = *N* (See Appendix D). Since the average response to mixtures is monotonically increasing with the rank correlation between *p* and 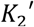 (See Appendix D), it is a simple corollary that 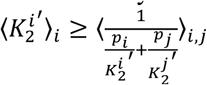 if *p* is perfectly anti-rank-correlated with 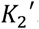. However, this also means that it is possible that 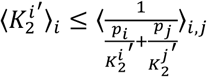 if the rank correlation between *p* and 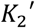 is sufficiently large. Therefore, while rough equivalence in average response to single odours and mixtures is observed for parameter choices corresponding to honey bees’ olfactory system, we cannot conclude that it is a general property of the receptor model.

**Figure 7:**
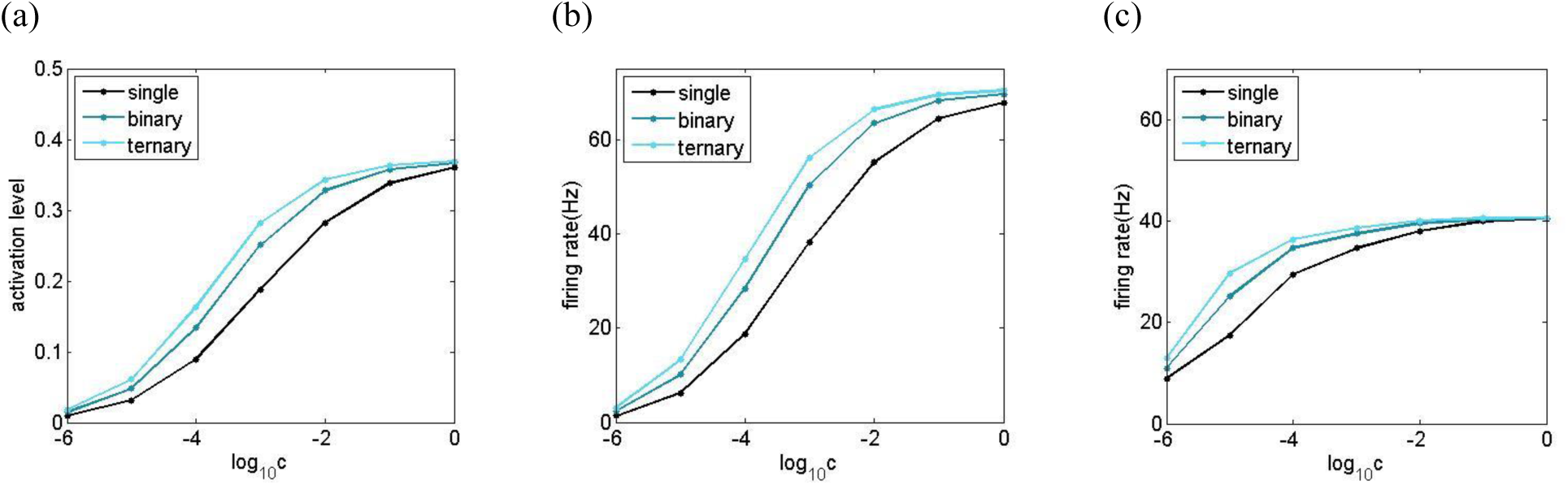
The relationships between the stimulus concentration and the average response for (a) ORN activation level (b) ORN firing rate and (c) PN firing rate, across all different odour-unit combinations. For both the activation level and firing rate, the average response strength for binary and ternary mixtures are larger than those of components but smaller than twice and triple those of single components at low stimulus concentrations. They, however, become almost identical at high stimulus concentrations. Note that the results for ORNs are obtained by further averaging over 1000 trials. Error bars are not shown as the variability of the results across trials is very small (*σ*firing rate < 0.6, *σ*activation < 0.003), in particular at high stimulus concentrations.

#### 3.2.3 Correlation between asymptotic response patterns at high and low stimulus concentration

Figure 8 shows that the asymptotic ORN response patterns to the same single component odour stimulus at high and low concentration are less correlated than those of mixture stimuli, suggesting ORN response patterns to mixtures tends to be more concentration-invariant than those of single component odours.

**Figure 8:**
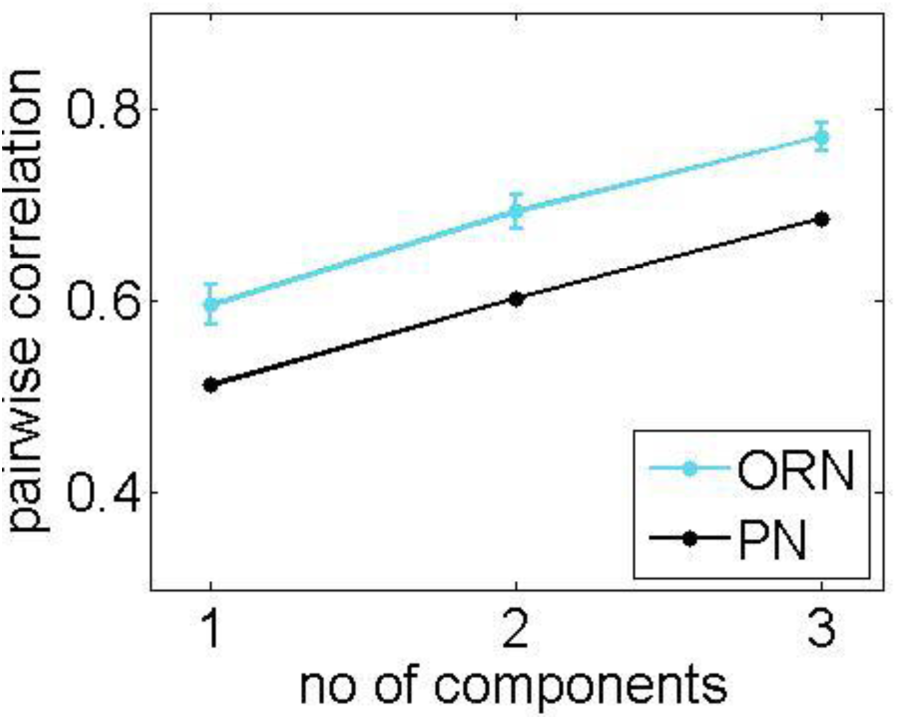
The pairwise correlation, averaged over all ORNs and PNs, between the response patterns at low (c = 10-^4^) and high (c = 10-^1^) concentration. For ORNs, the results is an average of over 1000 trials and the error bar is the standard deviation across different trials. We also verified that the observed monotonic relationship between the cross-concentration correlation and the number of components holds for every single trial. For PNs, the results are from a single trial.

In section 3.2.1, we have shown that the activation levels of receptors at the limit of low and high effective stimulus concentration are K_eff_ c_eff_ and 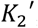 (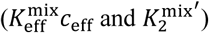) for odours of a single (multiple) component(s). Therefore, the correlation between activation patterns for stimuli in the limit of low and high ceff is essentially equivalent to the correlation of K_eff_ and 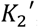, assuming that ceff for each odour-receptor combination is the same. If K_eff_ and 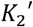, are strongly positively correlated, weak response at low concentration is more likely accompanied with weak response in high concentration for that odour-receptor combination, and vice versa. Taking the case of a single component odorant as an example, we illustratively show in Figure 9 that there is a strong positive dependence of the correlation between response patterns at different effective concentrations and K_eff_ and 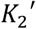.

**Figure 9:**
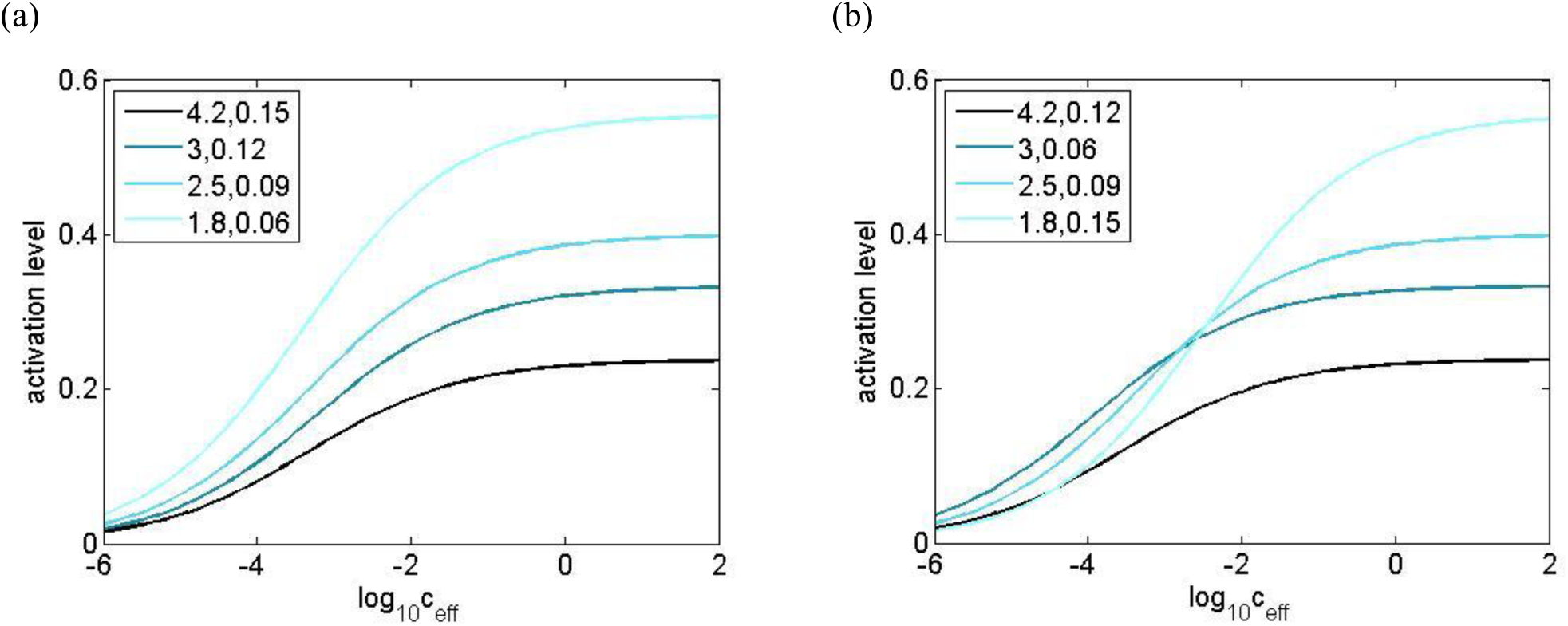
Dose response curves for different K_eff_ and 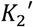, where the correlation between the parameters are (a) strongly positive and (b) non-positive. When K_eff_ is strongly positively correlated with 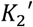, the proportion between the responses of different units are roughly constant over a large range of effective concentration, which indicates a high linear correlation between the response patterns at different effective concentrations. The opposite is observed if they are not strongly positively correlated. Legend format: first number 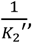, second number 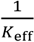

If we assume that k1, k-1 and K2 are independent, the pairwise Pearson correlation between K_eff_ and 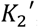 is positive as a simple consequence of the Chebyshev integral inequality (See e.g. Egozcue et al, 2009) since K_eff_ is monotonically increasing with K2 and K2 is monotonically increasing with 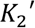.

We then asked, whether, under these assumptions, 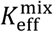 and 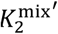 would be more strongly positively correlated than *K eff* and 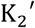 While it is not expected that the above statement holds universally, we computed the correlations between 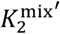 and 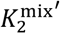, and K eff and 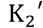 using a number of parameter sets with different ranges and statistical distributions, including biologically plausible ones, over many trials. We were able to verify that the above statements hold true for all trials even if the distribution of K1 and K2 are strongly skewed, as shown in Table 2. Therefore, we are confident that the above-mentioned conjecture holds for most, if not all, reasonable probability distributions of K1 and K2.

**Table 2:** The difference in the mean correlation between K_eff_ and 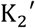 and that of 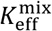 and 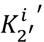, and the cross-eff 2 eff 2 concentration (c = 10-^4^ and *c* = 10-^1^) correlation between the response patterns, in terms of firing rate, to binary mixtures and single component odour stimuli over 1000 trials. The correlation of 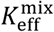 and 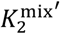 is higher for all trials and for all choices of parameter sets. The variability of the “transduction constant” *n* (n’: log-normal distribution, 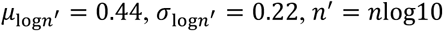, chosen based of experimental measurements by Gremiaux et al (2012)) weakens the effects and introduces discordance in some of the trials. However, the cross-concentration correlation of the response patterns for mixtures is still significantly higher than that of single component odour stimuli and instances of discordance are rare. *A hard boundary of K1, K2 > 0 is imposed for unbounded distributions. ** Non-biologically plausible parameter sets

| Probability distribution | $k_1^n$ (min,max)/ $\mu, \sigma$ | $k_{-1}$ (min,max)/ $\mu, \sigma$ | $K_2$ (min,max) / $\mu, \sigma$ | Mean corr difference | % of discordant trials |
| --- | --- | --- | --- | --- | --- |
| **$K_{\text{eff}}^{\text{mix}}$ and $K_2^{\text{mix}'}$, and $K_{\text{eff}}$ and $K_2'$ ($n = 0.65$)** |  |  |  |  |  |
| Uniform | (0.5,5) | (0.005,0.05) | (0.01,1) | 0.061 | 0 |
| Exp(uniform) | (0.63,31.6) | (0.006,0.1) | (0.01,1) | 0.095 | 0 |
| Normal * | 4,1.5 | 0.03,0.01 | 0.3,0.15 | 0.038 | 0 |
| Uniform** | (0.5,5) | (0.005,0.05) | (1,10) | 0.06 | 0 |
| Uniform** | (0.01,0.1) | (0.1,1) | (0.01,1) | 0.061 | 0 |
| Exp(uniform)** | (0.01,1) | (0.01,1) | (0.01,10) | 0.063 | 0 |
| Log(uniform)** | (0.095,4.61) | (0.001,0.095) | (0.01,1.1) | 0.042 | 0 |
| **Average firing rate for mixture and single component odourant ($n = 0.65$)** |  |  |  |  |  |
| Uniform | (0.5,5) | (0.005,0.05) | (0.01,1) | 0.239 | 0 |
| Exp(uniform) | (0.63,31.6) | (0.006,0.1) | (0.01,1) | 0.379 | 0 |
| Normal * | 4,1.5 | 0.03,0.01 | 0.3,0.15 | 0.312 | 0 |
| **$K_{\text{eff}}^{\text{mix}}$ and $K_2^{\text{mix}'}$, and $K_{\text{eff}}$ and $K_2'$ (variable $n$)** |  |  |  |  |  |
| Uniform | (0.5,5) | (0.005,0.05) | (0.01,1) | 0.056 | 0 |
| Exp(uniform) | (0.63,31.6) | (0.006,0.1) | (0.01,1) | 0.096 | 0 |
| Normal * | 4,1.5 | 0.03,0.01 | 0.3,0.15 | 0.029 | 0 |
| **Average firing rate for mixture and single component odourant (variable $n$)** |  |  |  |  |  |
| Uniform | (0.5,5) | (0.005,0.05) | (0.01,1) | 0.083 | 9 |
| Exp(uniform) | (0.63,31.6) | (0.006,0.1) | (0.01,1) | 0.308 | 0 |
| Normal * | 4,1.5 | 0.03,0.01 | 0.3,0.15 | 0.101 | 7 |

The remaining questions are whether the higher correlation between 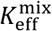 and 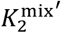 can be extended to correlation of response patterns, in terms of firing rate, at different concentration, and whether such correlations would be destroyed by the non-uniformity of ceff for different odour-receptor combinations for stimuli with the same *c* caused by variability in n. Table 2 shows that, if the transduction coefficient *n* is a constant, the correlation of the firing rate response patterns at high and low ceff for binary mixtures is indeed higher than that of single component odours. This holds for all trials and for all tested biologically plausible parameter sets. If we also take into account the non-uniformity of *n* across ORNs, the difference in correlation decreases and it is possible that results from a few trials become discordant. However, Table 2 shows that overall the cross-correlation for mixtures is still significantly higher than for their single odours counterparts and the instances of discordance are very rare even when *n* is highly variable. This suggests that the higher cross-concentration correlation between response patterns is a general result for the receptor model as described in (3) and (4).

For non-biologically plausible parameter sets marked with ** in Table 2, the majority of the neurons either remain inactive at high concentration or already respond at the asymptotical level at low concentration. Correlation becomes non-indicative as a measure in such a context.

#### 3.2.4 Response latency

The response latency, defined as the time required for an ORN to fire a spike after stimulus onset, is primarily determined by the transient receptor response at the limit of small time. The stronger the transient response, the lower the latency. In Appendix F we consider the full sets of dynamics equation for single component stimuli and mixtures and find an approximation for r* and *r*_mix_* at the limit of small ceff and t. In the approximation, we assume that 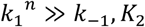, which is essential for both reproducing the rapid pulse tracking ability of ORNs observed in Szyszka et al (2014) (See also Figure 2) and ensuring a realistic magnitude of receptor responses at different concentrations. This leads to the following expressions:

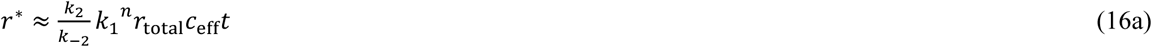

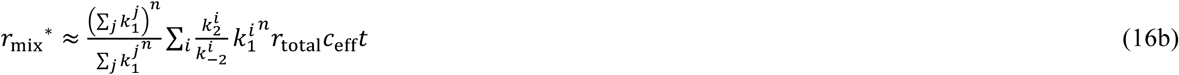

Now, we compare the average response to N-component mixtures and single component odorants for a typical receptor,i.e. we are averaging over *k* values, but not over n. To ensure that differences are not caused by the disparity in the number of molecules present in the single component odour and in the mixture, we consider the mixtures with concentration c0 for each component, and single component odorants with concentration Nc0. Using a similar approach as in the derivation of (15), we have for mixtures

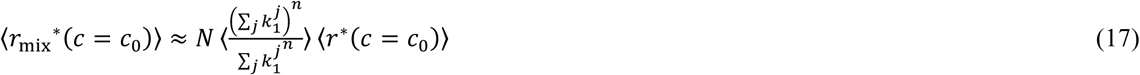

Note that the assumption of essentially uncorrelated 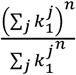 and 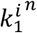 is made as in (14). For single component odorants with normalized concentration and assuming *n* < 1, we have

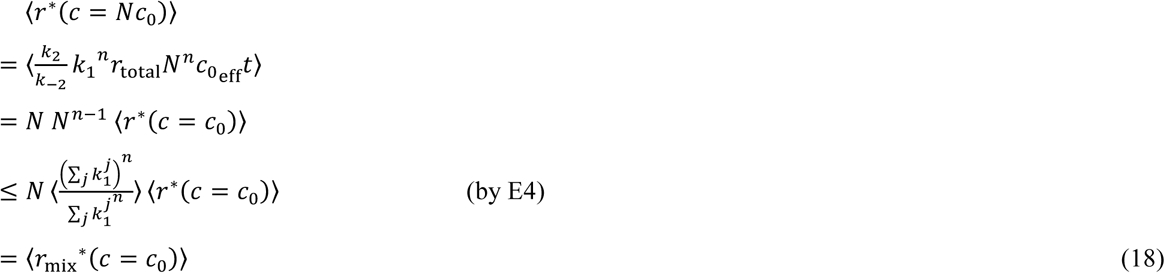

It can easily be shown that by (E3), the inequality sign in (18) flips if *n* ≥ 1.

As discussed in the previous section, *n* ≤ 1 for the majority of the receptors. This implies that on average one would expect that at the limit of small ceff, the response latency for mixtures is smaller than its single component counterpart with the same number of molecules, which has been verified by the results from simulation of our models as shown in Figure 10.

**Figure 10:**
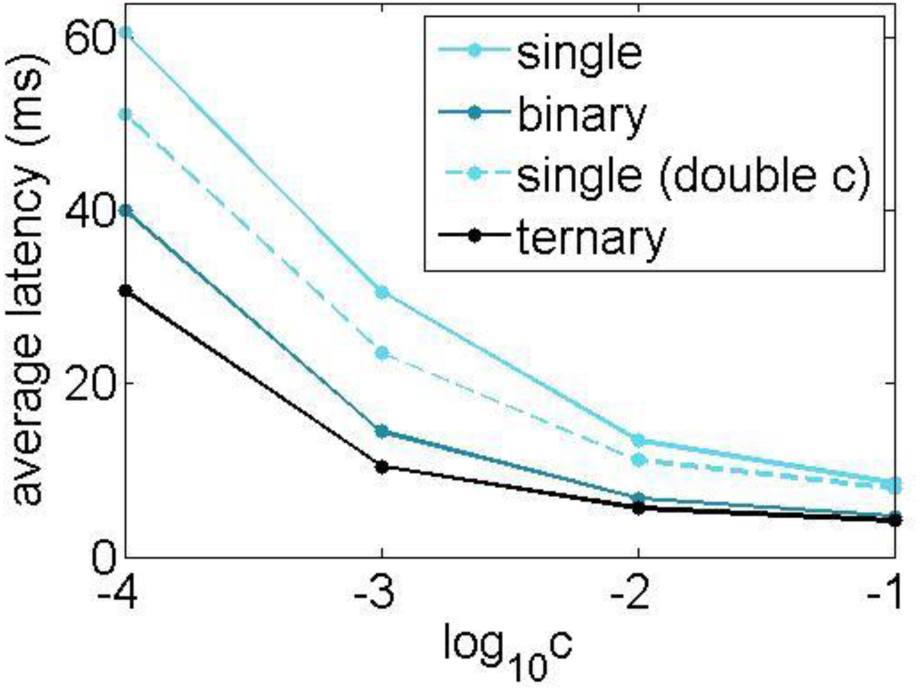
The average response latency decreases with the number of components in the odour stimulus. The effect is most significant when the stimulus concentration is low. This effect cannot be fully explained by the higher number of odour molecules in stimuli with more components compared to their counterparts with less components at the same concentration, since the latency for binary mixtures (red) is lower than that for a single component odour with doubled concentration (green). Please note that an absolute latency of 1ms is added to the latency generated by simulation to mimic the time required for the diffusion of odour molecules in the sensilla.

At high concentration, Free receptors are quickly depleted and equilibrium is quickly reached such that the approximation in (16a) and (16b) breaks down already at very small t, much earlier than the receptor neuron can fire a spike. In this regime, the response latency of the neurons is dominated by the equilibrium activation level, which on average, as shown in Figure 1, is almost equivalent for single component stimuli and mixtures. This offers an explanation why the average response latencies for these two type of stimuli are also almost equivalent at high concentration.

#### 3.2.5 Monotonicity of the results with respect to the number of components in mixtures

Although the monotonicity of the above results can be established by further simulation and analysis, one can obtain an intuitive understanding of it by considering the following: We can interpret a ternary mixture as a binary mixture of a binary mixture of two components with the third component. We can then consider the binary mixture as a single odour by transforming 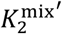 and 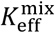 in (11b) into 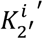 and 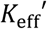, and apply the analyses in the previous sections 2 eff 2’ eff with the third component being the second odour in the mixture (taking the value of 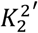 and 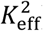). This procedure 2 eff can be repeated to obtain results for mixtures having an arbitrary number of components, and the validity of this approach is shown in (C15). Following this idea, one can clearly see that the any change in response properties with respect to the single component case must be monotonic as the number of components in the mixtures increases.

#### 3.2.6 Mixtures of different proportions

If the concentration of the components in a mixture is not identical, we can add weighting terms to the terms in the summation in 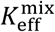 and 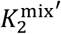, so that 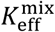 becomes a weighted sum of 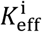 while the weight *p*_i_ in 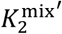 is further weighted by the effective concentration for different components (See e.g. (C11)). It can be expected that much of the discussion above would not be affected as these weighting effects would be averaged out, except that the magnitude of change for different properties may be altered. The detailed study for such effects is beyond the scope of this work.

#### 3.2.7 Extension of the results to projection neurons

One may ask whether our observations at the level of receptors and receptor neurons would also be preserved in PNs. If we assume that spike generation is independent of the sub-threshold membrane potential dynamics in the ORNs, it is obvious that lower 1st spike response latency in the ORNs implies lower latency in PNs. For the observations of weaker response gain and stronger correlation between response to stimuli at low and high concentrations in ORNs,one might think that the non-linearity of spike generation and inhibition may render the observed effects insignificant in PNs. We show in Figure 7c and 8 that this is not the case.

## 4. Discussion

In this work, we produced a biophysical model of the early olfactory system of honey bees in which their full receptor repertoire is considered. Instead of directly fitting specific experimental data, model responses are generated with constraints from the overall statistics observed in ORN responses in a number of experiments, namely the generally high pairwise correlation between ORN response to different odours (Galizia et al, 1999), the dependence of similarity in ORN response pattern and chemical structure of odorants (Carcaud et al, 2012), the sigmoid-shaped response-dose relationship (Rospars et al, 2008, Gremiaux et al, 2012), and the fast response dynamics and high sensitivity to temporally fluctuating inputs (Syzszka et al, 2014). With this approach, we showed that many experimental findings from data not used to build the model can be reproduced.

The model takes into account several biophysical processes at a minimal level, including processes of chemical binding and activation in receptors, and spike generation and transmission in the antennal lobe network. This allows us to pinpoint which part of the system is responsible for certain features of the response. At the same time, it is simple enough to allow mathematical analysis and/or efficient numerical simulation of various parts of the system.

We then extended the receptor model such that it can also describe responses to mixtures of chemicals. The model can reproduce experimentally observed synergistic and hypoadditive mixture responses (Duchamp-Viret et al, 2003; Rospars et al, 2008; Cruz and Lowe et al, 2013). We then applied probability theory in order to compare the statistical properties of the receptor responses to single component and mixture stimuli, and showed that for binary mixtures the response latency of ORNs at low stimulus concentrations is reduced and the response patterns are less variable across concentrations, as compared to single component odours. We showed rigorously that these results are not specific to any particular fine-tuned parameter set, but are a general consequence of the receptor dynamics described by (4) in a large regime containing biologically plausible scenarios. In addition, our results imply that any change in response properties with respect to the single component case must be monotonic with the number of components in the mixtures. Finally, by numerical simulation, we found that these results are preserved in PNs, unaffected by processes in the later part of the system, including spike generation and LN inhibition.

### 4.1 Improvements of our receptor model for mixtures over previous models

In our receptor model for mixtures, the total rate of molecule binding is determined by applying the transduction cascade after summing up the effective affinity to the receptors kjcj for each component. Previous work (Rospars et al, 2008; Nowotny et al, 2013) instead applied the transduction cascade to each of the components, which gives equations of the following form:

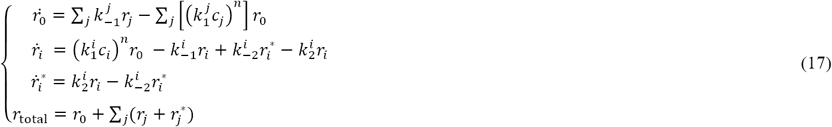

Cruz and Lowe (2013) pointed out that this equation is inconsistent for‘mixtures’ of identical components with randomly partitioned concentrations. They offered a solution by not considering the transduction constant *n* in the dynamical equations but added them directly to the solution. It is highly non-trivial to ascertain whether such altered expressions would correspond to the solutions to a system of equations that have good biological basis. Our approach is more transparent on how molecules may interact biophysically, and offers experimentally testable predictions.

In our model, the rate of receptor binding for each component is weighted by its effective affinity to the receptor after taking transduction into account. We believe this is the best educated guess based on the information we have. We assumed that the total rate of receptor binding depends on transduction processes, which means such processes influence the interaction between odour molecules and receptor sites, and competition between different types of molecules for receptor sites should also take into account such effects.

### 4.2 Limitations of our model

In the model, we did not consider inhibitory responses for ORNs. A technical reason is that experimental measurements using calcium imaging are not very good in exposing inhibitory responses. While some ORNs do express inhibitory responses (Getz and Akers, 1993; de Bruyne et al, 1999), it is unlikely that they contribute substantially to odour identification, since the system would otherwise need to take much longer to perform odour identification than results from behavioural experiments suggest (Resulaj et al, 2015) given the relatively small number of neurons in the bees’ early olfactory system. It is possible though that inhibitory responses are important for coding fine features of odorants but that is beyond the scope of this study. With suitable experimental data the model can be adjusted by changing a few parameters to also generate inhibitory responses.

We have made several approximations when calculating the firing rate and response latency of neurons. The instantaneous firing rate v, by our definition in (10), corresponds to the anticipated firing rate if the effective input current *I*_eff_ (see (8)) stays constant. The lack of a temporal filter representing the finite time scale of spike generation leads to an overestimation in the sensitivity of neurons to input fluctuations (Ostojic and Brunel, 2011; Schaffer et al, 2013). However, in this context, such effects are alleviated by the additional temporal filters in the process of binding and activation of receptors, as described in (3), which smoothen out *I*_eff_ to some extent even if the stimulus intensity fluctuates rapidly. Moreover, we assumed that inputs are noise-free when calculating the firing rate and response latency of ORNs. This assumption is valid for dominantly mean-driven neurons firing regularly at high rate. Outside this regime, the firing rate and response latency of the neurons are underestimated. However, as we have explained in the previous paragraph, these slow-firing neurons are unlikely to be essential for odour identification, and given the low spontaneous firing rate observed in most ORNs, it is unlikely that noise would affect the AL response substantially apart from cases when the stimulus is very weak. Under certain circumstances, it is possible to approximate higher-dimensional conductance-based models by a 1-dimensional integrate-and-fire model (Fourcaud and Brunel, 2002; Richardson and Gerstner, 2005). In such cases, a good approximation can be obtained for the mean firing rate and the probability distribution of response latency for arbitrary input without having to directly simulate (7) numerically (Ostojic and Brunel, 2011; Chan et al, in preparation). These methods also do not require an adiabatic approximation.

To generate the ORN responses, we assumed that all the interactions take place at the receptor level. ORNs would integrate input from receptors but do not interact with each other. Experimental measurements (Duchamp-Viret et al, 2003; Rospars et al, 2008) have shown that for a small minority of odour-receptor combinations, ORN responses can decrease with concentration. In addition, there is a small chance that mixture responses are lower than all the individual responses to their constituent components. While this can be reproduced by having the transduction constant *n* taking a negative value, it is unrealistic because of the singularity at zero concentration. A more plausible mechanism for such results is coupling between different ORNs. It has been reported in *Drosophila* that excitation of a receptor neuron may inhibit its neighbour in the sensillum via ephaptic interactions (Su et al, 2012; Van der Goes van Naters, 2013), which may lead to the above-mentioned suppressed mixture responses. However, since the structure of sensilla in bees’ antenna is different from that of *Drosophila* sensilla, it is unclear whether the same effects would exist in bees. Better knowledge about the bees’ anatomy and chemical processes in the receptor is required to improve the model. Nevertheless, here we have shown that even without considering the above-mentioned extra interactions, many features of olfactory processes can already be reproduced. This raises the question whether and how these interactions affect olfactory processing and coding.

### 4.3 The role of the 2-step binding and activation processes in olfactory receptors

It is well known that there are two distinct chemical processes taking place in olfactory receptors: binding of odour molecules to receptors and activation of bound receptors (Rospars et al, 2008). One may ask whether a single process would be sufficient to produce olfactory responses without compromising the coding of stimuli. Our results ((11a) and (11b)) elucidate how the olfactory response depends on each process. For single component stimuli, the asymptotic response in the limit of high concentration depends only on the activation process while the response at the other limit of low concentration depends on both the binding and activation processes. From this, it is clear that the process of activation of bound receptors is essential for a rich representation of different stimuli at high concentration, and thus identification of odour in this regime. However, it also somewhat destroys the correlation between the representations for the same odour at different concentrations. Here we showed that this can be alleviated by replacing single component odorants with mixtures, since the response to mixtures at high concentration no longer only depends on the activation of receptors but also is weighed by the affinity of the constituent odorants to the receptors. The additional dependence on the binding step leads to the higher correlation between response patterns at high and low concentration.

### 4.4 Implication of our results on coding of odours

An important question is how the lower average response latency and higher correlation between responses across concentrations for mixtures affect information coding. Behavioural experiments (Resulaj et al, 2015) suggest that odour identification by the olfactory system is performed in the time scale of 101ms. Here we describe two simplistic code which is possible under such constrains. First, the response patterns are sampled for a fixed amount of time, after which a decision of odour identity is made (Junek et al, 2010; Wilson et al, 2017). With lower latency, more ORNs can be recruited for the identification of a particular odour. This implies larger information capacity of the system for mixture stimuli. Second, the decision of odour identity is determined by the response of a fixed number of ORNs (). In this case, the lower average latency for mixtures allows them to be identified by the system more quickly.

In natural environment, odours molecules are primarily transported through air by convection. Previous studies (Murlis et al, 1992; Weissburg et al, 2012) showed that odours molecules move through air in filaments, forming complex odour plumes. Very often, the movement of those plumes is chaotic because of turbulent air flow, which results in rapid and unpredictable fluctuations in the concentration of odours as received by animals. Therefore, to identify an odour, the response of the olfactory system need to be reasonably robust against changes in odour concentration. The strong positive correlation between response patterns across concentrations for mixtures is therefore highly conducive to odour identification.

One may argue that such correlations hinder the coding of odour concentration. There are several alternatives of how concentration information can be coded, as discussed in previous work. For instance, information for concentration may be coded by other features of the response, like the portion of activated glomeruli (Stopfer et al, 2003; Strauch et al, 2012), the connectivity between units with plastic synapses (Hopfield, 1991), or by responses at the later part of the olfactory system (Froese et al, 2013). It is also possible that concentration information is encoded by the temporal patterns of input, for instance the response latency of all or a sub-set of units (Hopfield, 1995) or degree of synchrony between firing of different units (Brill et al, 2015). Therefore, the improved identity coding due to more invariant asymptotic response pattern does not necessarily imply a compromise in concentration coding, and it remains an open question how identity and concentration coding may be simultaneously achieved (Hopfield, 1991,1995; Stopfer et al, 2003, Arnson and Holy, 2013).

## Acknowledgments

This work was partially funded by the Human Frontiers Science Program (grant RGP0053/2015) and the Engineering and Physical Science Research Council of the UK (grant EP/P006094/1).

## Appendix A Generating ORN responses which matches the statistics of experimental data

The method we used to generate the response vectors is inspired by that used by Haenicke (2015). The response vector of the *k^th^* ORN is denoted as *u_k_*, in which each element corresponds to the response of the ORN to a particular odour. For *k* ≤ 26, *u*_k_ was directly adopted from experimental data of the 26 ORNs as mentioned in the last paragraph. The rest of *u*_k_ were generated as follows:

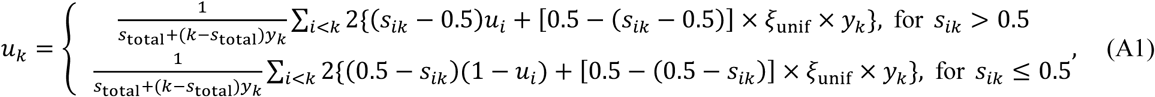

where *s*_*ij*_ are elements of an upper triangular matrix *s* which describes the ratio of correlation between pairs of glomeruli response vectors, *y_j_* controls the overall level of correlation between the response vector of the *j^th^* ORN unif and that of the other ORNs, *ξ*_unif_ is a uniform random number in [0,1). The factor 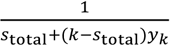, in which 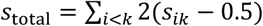, normalizes the responses to [0,1].

The statistics of the elements in each response vectors *r*_k_ (k > 26) generated by the above procedures does not follow that of their counterparts in the experimental data (k ≤ 26). Therefore, we rescaled all the elements *u*jk, where j = 1,2,… 16 corresponds to each of the 16 odours, in all response vectors *u_k_* for *k* > 26 and obtained new responses vectors 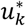 comprising of elements 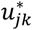, which are described by:

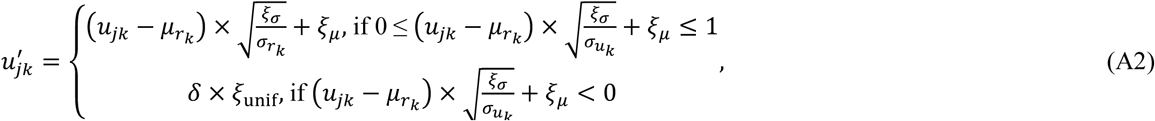

where *ξ*_*μ*_ and *ξ*_*σ*_ are random variables centred around the mean and the variance across all elements in all response vectors from the experimental data,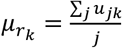, which corresponds to the mean of the elements in *u*_k_, 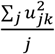, which corresponds to the variance of the elements in *u_k_*, δ is a small constant. There is a small j *k* possibility that the rescaled elements have negative values. In such cases, the rescaled value were artificially set to a value close to 0 as described by (A2). The rescaled elements were allowed to have value greater than 1 since it is possible that some ORNs which are not experimentally measured respond more strongly to some odours than all of its experimentally measured counterparts.

Both the statistics of the pairwise correlation of the generated vectors 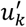 as well as the mean and variance across their elements match that of the experimental data. For the rest of the paper, *u_k_* refers to 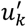 as described by (A2) instead of its counterpart in (A1).

Table A1: listed the parameters used for the random variables. They are obtained by manual trial-and-error. While our method and parameters provide generated response patterns with an excellent fit with the data, there may be other possible parameters or methods which may produce equally good or better fit with the data, the study of which is outside the scope of this work.

**Table A1.** Parameters used for (A1) and (A2). *A hard boundary 0 < *s* < 1 is imposed. **A hard boundary 0 < *n* < 2 is imposed.

| Random variables | pdf | $\mu, \sigma/(\text{min, max})$ |
| --- | --- | --- |
| $s$ | *mixture of 3 normal distributions | 0.88, 0.075 with $p = 0.2$<br>0.16, 0.075 with $p = 0.2$<br>0.64, 0.075 with $p = 0.6$ |
| $y$ | **normal | 1, 1 |
| $\xi_\mu$ | mixture of 2 uniform distributions | (0.12, 0.4) with $p = 0.75$<br>(0.35, 0.7) with $p = 0.25$ |
| $\xi_\sigma$ | Folded normal | 0, 0.028 |

### Appendix B: Parameters of the model

**Table B1.** Parameters used in the model.

| Random variables | | p.d.f. | value/$\mu, \sigma$ | units | remarks |
| --- | --- | --- | --- | --- | --- |
| **Parameters for the Hill curves in (2) and receptor dynamics** |  |  |  |  |  |
| $C_{\frac{1}{2}}$ | | normal | -3, 1 | | hard boundary: $-4.4 < C_{\frac{1}{2}} < -0.4$ |
| $n'$ | | log-normal | 0.45, 0.3 | | hard boundary: $0.7 < n' < 3.5$ |
| $r_0$ | | constant | 2 | | |
| $k_1$ | | normal and then scaled* | 1.2, 0.15 | ms<sup>-1</sup> | scaling factor: $\sqrt{C_{\frac{1}{2}} n'}$<br>hard boundary: $0.1 < k_1 < 5000$ |
| $k_2$ | | normal | 0.1, 0.01 | ms<sup>-1</sup> | hard boundary: $k_2 > 0$ |
| Note: $k_{-1}$ and $k_{-2}$ are constrained by the above. Hard boundary: $k_{-1} > 0.01, 0 < k_{-2} < 50$ | | | | | |
| **AL network connectivity between units**<br>(Note in all cases a hard boundary of mean $\pm 2$ standard deviation are applied) | | | | | |
| $w_0$ | | normal | 0.006, 0.002 | | |
| $w_{\text{corr}}$ | | normal | 0.01, 0.001 | | |
| $g_{\text{ORN}}$ | | constant | 2 | nS | |
| ORN-PN |  | normal | 0.045, 0.01 |  |  |
| ORN-LN |  | normal | 0.013, 0.003 |  |  |
| LN-PN |  | normal | 0.04, 0.01 |  |  |
| LN-LN |  | normal | 0.004, 0.001 |  |  |
| **Spiking model** |  |  |  |  |  |
| $\tau_m$ | | constant | 20 | ms | |
| $V_e$ | | constant | 50 | mV | |
| $V_i$ | | constant | -75 | mV | |
| $V_r$ | | constant | -70 | mV | |
| $g_l$ | | constant | 1 | nS | |
| $V_{th}$ | | constant | -50 | mV | |
| $V_{\text{reset}}$ | | constant | -70 | mV | |
| $t_{\text{refract}}$ | | constant | 2 | ms | |
| background excitation | ORN | constant | 0.28 | nS |  |
| | PN | normal | 0.24, 0.02 | nS | hard boundary: mean $\pm 2$ sd |
| background inhibition | ORN | constant | 0.5 | nS |  |
| | PN | normal | 0.15, 0.01 | nS | hard boundary: mean $\pm 2$ sd |
| $I_{\text{adapt}}^{\text{base}}$ | ORN | constant | 40 | mA | |
| | PN | normal | 4.5, 0.4 | mA | hard boundary: mean $\pm 2$ sd |
| | LN | normal | 1.8, 0.2 | mA | hard boundary: mean $\pm 2$ sd |
| $\tau_{\text{adapt}}$ | ORN | constant | 60 | ms | |
|  | PN/LN | constant | 25 | ms |  |

### Appendix C: Derivation of the asymptotic receptor response

Here we are deriving the asymptotic response for the dynamical equations (3) and (4). The dynamics equation of binding and activation for single odour stimulus is

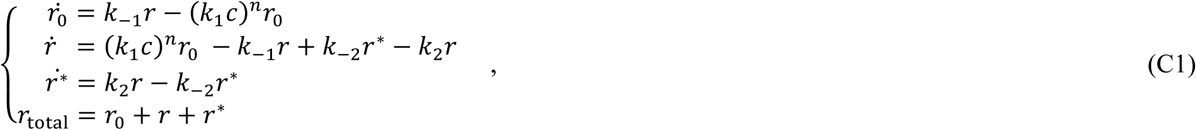

At equilibrium, we have 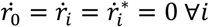, this gives

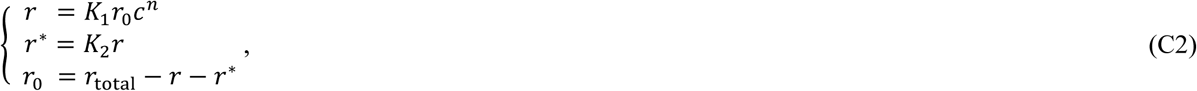

where 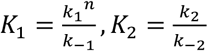

Hence we have

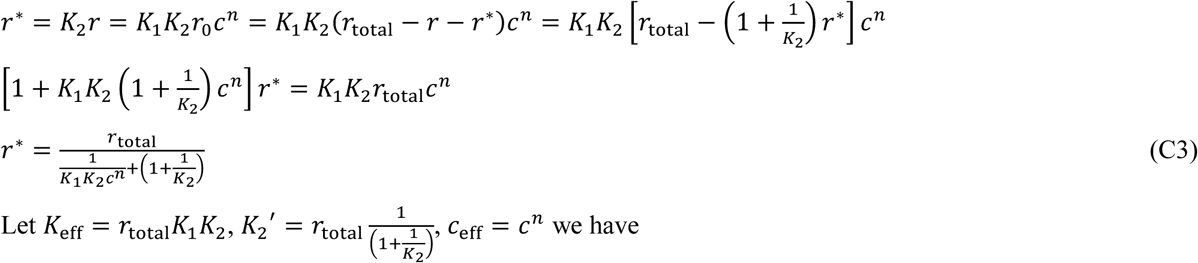

Let *K*_eff_ = *r*_*total*_*k*_1_*k*_2_,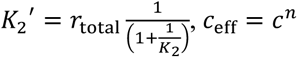,*ceff* = cn we have

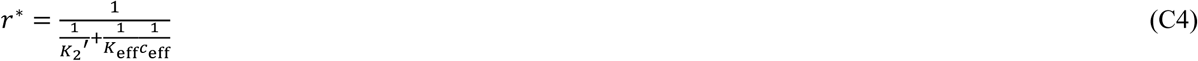

We now consider, more generally, the sets of dynamics equation for stimulus with an arbitrary number of components:

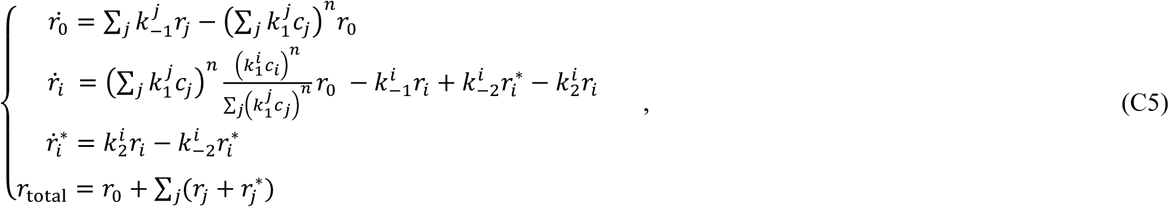

where the subscript i refers to the parameter for the ith component in the stimulus At equilibrium, we have 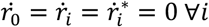, this gives

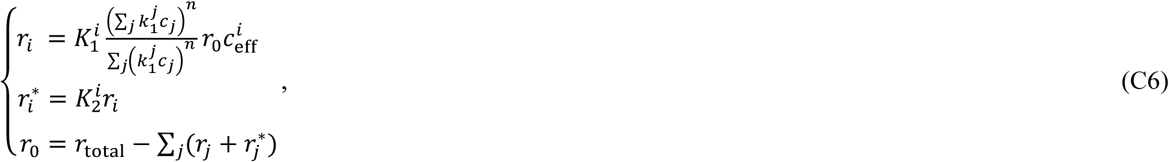

Hence we have

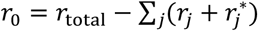

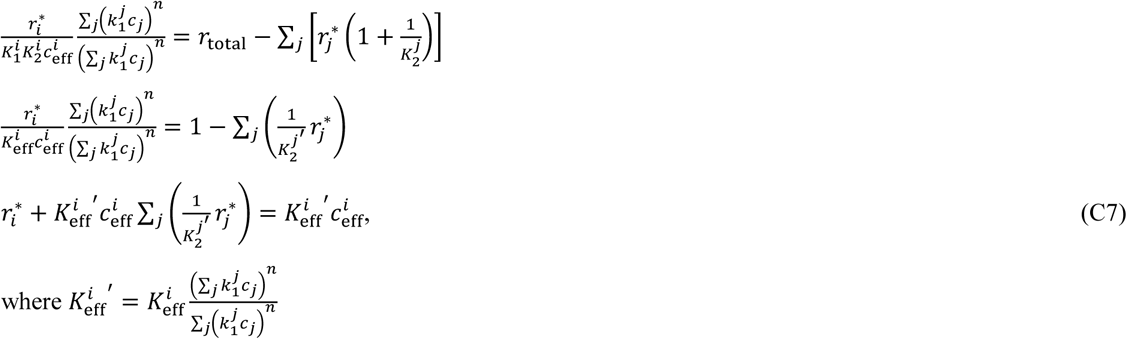

Where 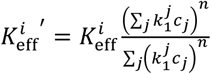 (C7) can be rewritten in matrix form,

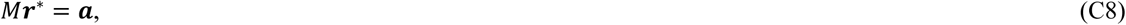

Where 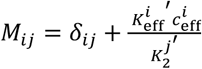 is the Kronecker delta function,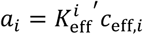 r* can be obtained by finding the inverse of M, and the total receptor response 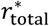 is given by 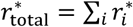 First we will look at the special case of binary mixture, in this case 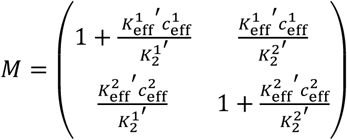

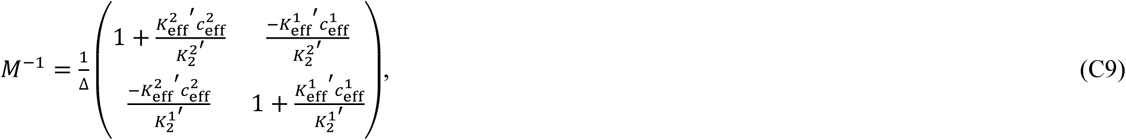

Where 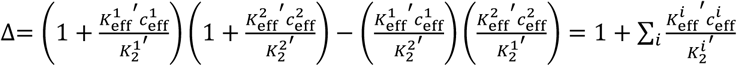

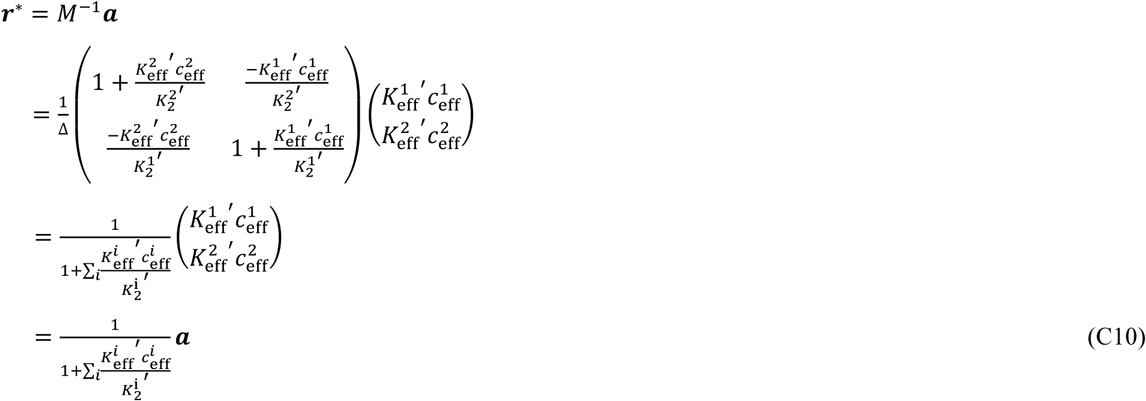

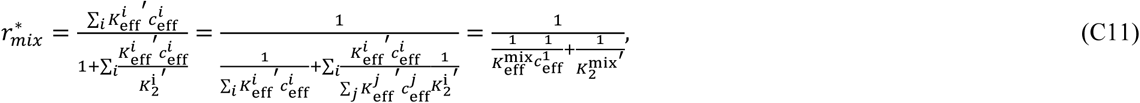

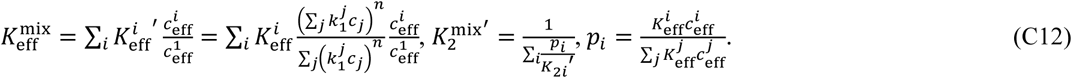

If 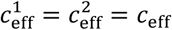 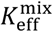 and *p*i can be further simplified as

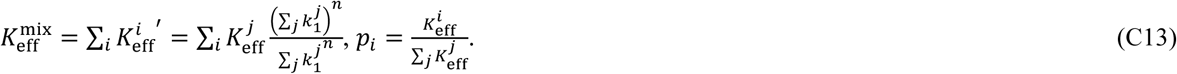

In fact (C10) to (C13) is true for mixture with any number of component, note that

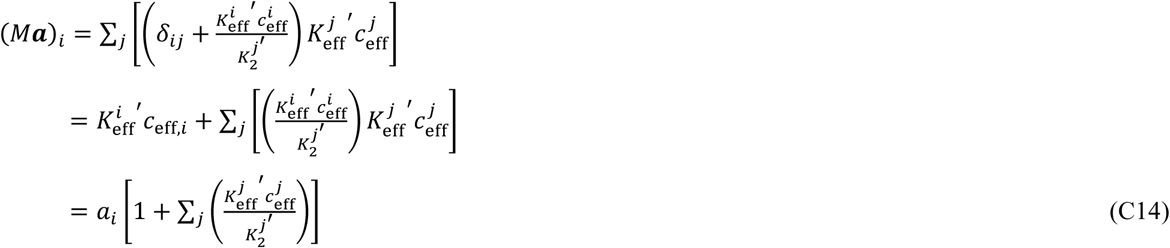

We have 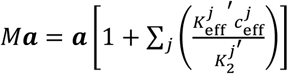, which gives (C10)

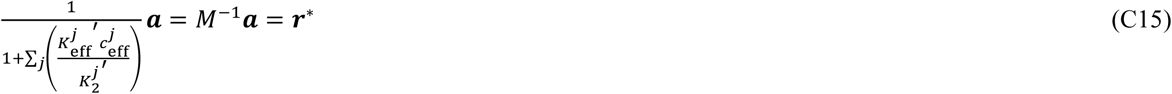

### **Appendix D: A proof of weaker average receptor response to binary mixture than single odour in the limit of high stimulus concentration, assuming homogeneous** *K*_*eff*_

We are now proving where 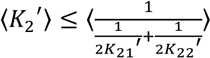 and 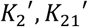 *and* 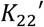 are uniformly and independently sampled from the set *Y* = {*y*1, *y*2,…, *y*N}

Let 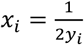 and arrange the terms such that x_1_ ≥ x_2_ ≥… ≥ x_N_ > 0

We want to show that 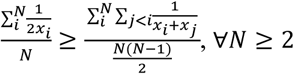

**Proof by induction on** *N*:

For *N*=2,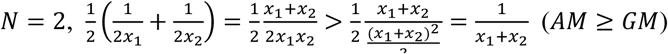

Assume it is true that 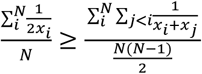, where x_1_ ≥ x_2_ ≥… ≥ x_N_ > 0

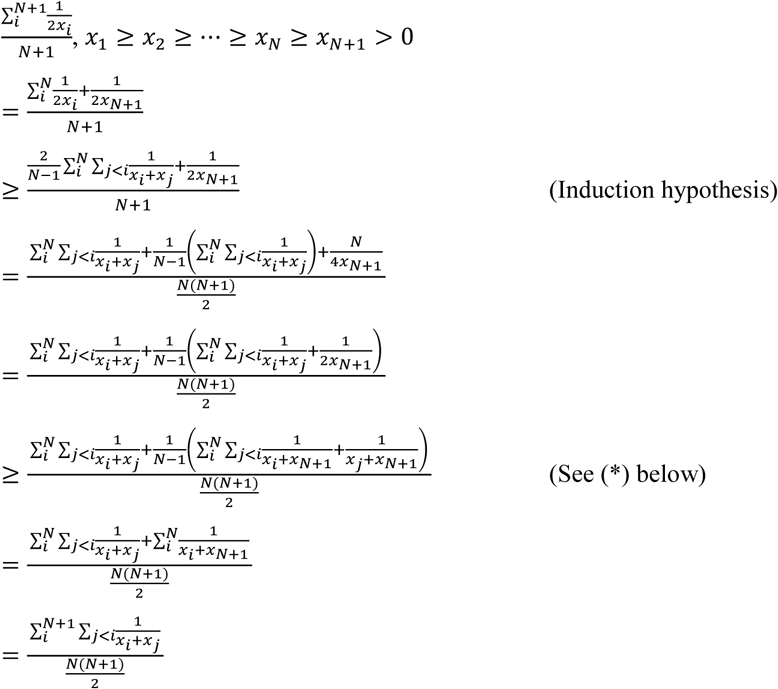

q.e.d.

(*)

To show 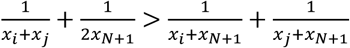

Note that 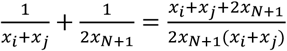 and 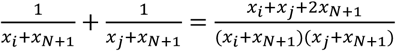

Since the numerator for both expressions are the same, by comparing the denominator, we have

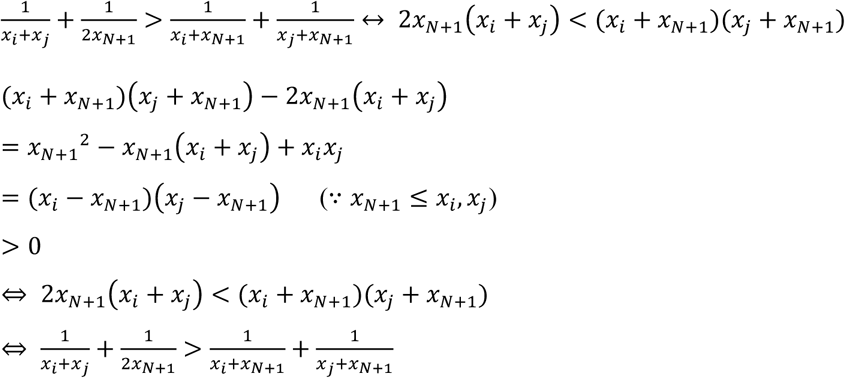

q.e.d.

To show 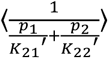 is increasing with its rank correlation between *p* and 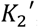, it is sufficient to show that 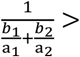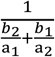 if a_1_ > a_2_ > 0 and b_1_ > b_2_ > 0. Note that

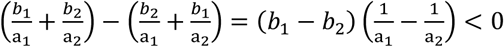

Therefore,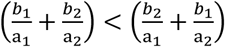, which implies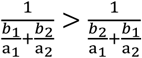.

### Appendix E

Here we provide a proof for (13a) and (13b).

Since the parameters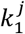 are strictly positive, we can rewrite 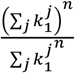 in terms of the *Lp*-norms of a vector k_1_ = 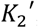,

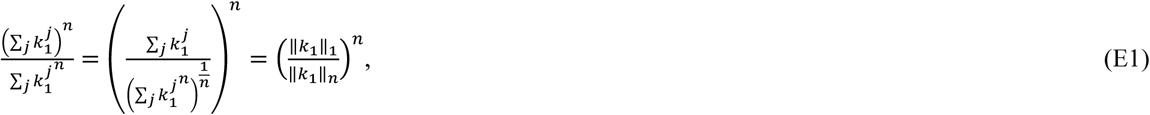

where ‖ · ‖_*p*_ corresponds to the *Lp* norm.

It is straightforward to show, e.g. by Janssen’s inequality, that ‖ · ‖_*p*_ decreases with *p*, ‖k_1_‖_n_ = ‖k_1_‖_1_ if and only if *n* ≥ 1. This gives

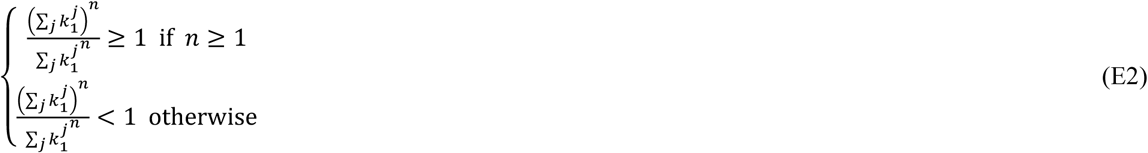

To obtain the upper (lower) bound for *n* ≥ 1 (n < 1), we use Hölder’s inequality. For *n* ≥ 1, we have

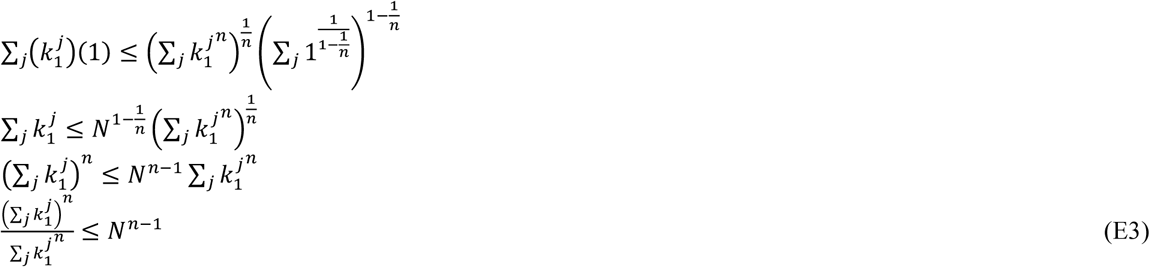

For *n* < 1, we have

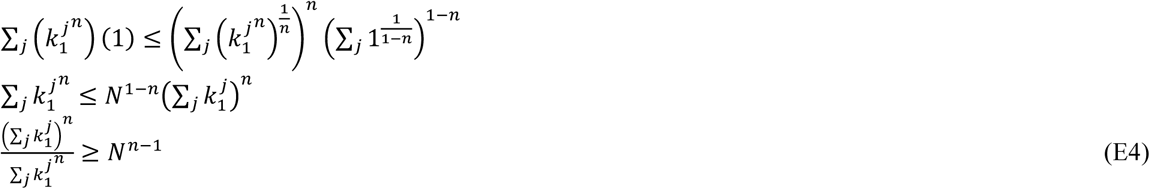

In *3.2.1*, we stated that it is possible to obtain suppressive response when *n* < -1. We illustrate this with the example of *n* = -2 using the Cauchy-Schwarz inequality.

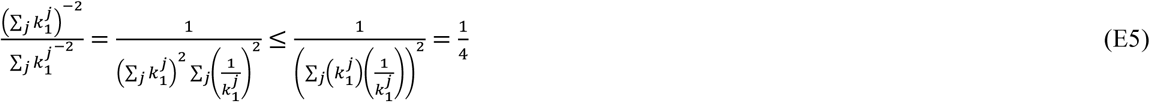

By (C13) and (E5),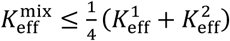, which can be smaller than both 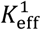 and 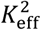

### Appendix F: Derivation for the receptor dynamics in the limit of small t

The dynamics equation of binding and activation for single odour stimulus is given by (C1)

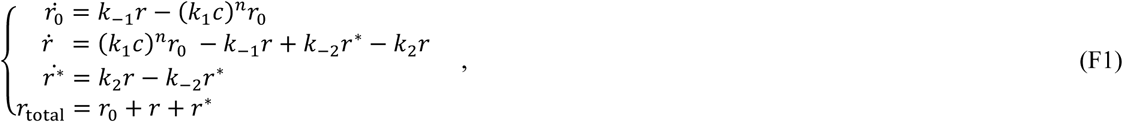

At small *t* and with the assumption that 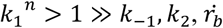 can be approximated by 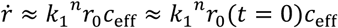 Since *r*_0_(t = 0)= rtotal and r(t = 0) = 0, we have

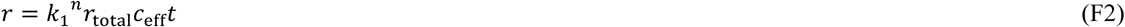

Substituting (F2) into the third equation of (F1) gives

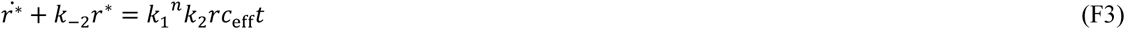

(F3) is linear and can be solved by standard methods. The solution is given below

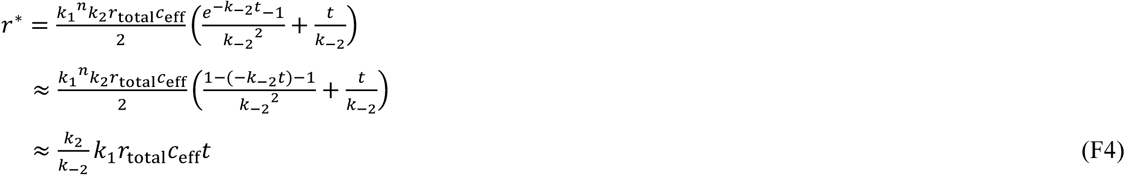

By comparing (E2) and (E4), we can see that the rate of increase in activation is slower than that of binding by a factor of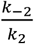 It is also clear that the approximation breaks down when *k*_2_ >_*k*-2_.

Similar calculation can be done for mixture stimulus. As the competition for binding sites for different odour components is negligible at small t, the binding and activation step is independent for each odour component. It can easily be shown that

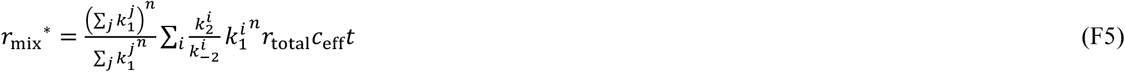

